# Regulation of hematopoietic stem cell (HSC) proliferation by Epithelial Growth Factor Like-7 (EGFL7)

**DOI:** 10.1101/2025.01.21.634107

**Authors:** Rohan Kulkarni, Chinmayee Goda, Alexander Rudich, Malith Karunasiri, Amog P Urs, Yaphet Bustos, Ozlen Balcioglu, Wenjun Li, Sadie Chidester, Kyleigh A Rodgers, Elizabeth AR Garfinkle, Ami Patel, Katherine E Miller, Phillip G Popovich, Shannon Elf, Ramiro Garzon, Adrienne M Dorrance

**Author notes:** These authors contributed equally (Rohan Kulkarni, Chinmayee Goda). Corresponding author – Dr. Adrienne Dorrance., **Email:**. **Classification:** Major – Biological Sciences, Minor – Cell Biology.

## Abstract

Understanding the pathways regulating normal and malignant hematopoietic stem cell (HSC) biology is important for improving outcomes for patients with hematologic disorders. Epithelial Growth Factor Like-7 (EGFL7) is ∼30 kDa secreted protein that is highly expressed in adult HSCs. Using *Egfl7* genetic knock-out (*Egfl7* KO) mice and recombinant EGFL7 (rEGFL7) protein, we examined the role of Egfl7 in regulating normal hematopoiesis. We found that *Egfl7* KO mice had decreases in overall BM cellularity resulting in significant reduction in the number of hematopoietic stem and progenitor cells (HSPCs), which was due to dysregulation of normal cell-cycle progression along with a corresponding increase in quiescence. rEGFL7 treatment rescued our observed hematopoietic defects of *Egfl7* KO mice and enhanced HSC expansion after genotoxic stress such as 5-FU and irradiation. Furthermore, treatment of WT mice with recombinant EGFL7 (rEGFL7) protein expands functional HSCs evidenced by an increase in transplantation potential. Overall, our data demonstrates a role for EGFL7 in HSC expansion and survival and represents a potential strategy for improving transplant engraftment or recovering bone marrow function after stress.

## INTRODUCTION

Hematopoietic stem cells (HSCs) reside within the bone marrow (BM) and retain the capacity to proliferate, differentiate and self-renew to provide lifelong blood production in mammals (1, 2). HSCs mostly reside in the long bones of adult mammals where they are regulated by various intrinsic and extrinsic factors that govern fate decisions. HSCs in the BM are relatively inactive and maintain their quiescent phase when compared to differentiated cells (3, 4). However, in stress conditions, HSCs are activated and increase their self-renewal and proliferation (5, 6). A comprehensive understanding of the mechanisms controlling HSC activation, survival, self-renewal/proliferation, and differentiation of HSCs will allow for the manipulation of HSCs *in vitro* for their use in novel cellular therapies. Currently, limitations in the availability and expansion of HSCs that retain robust long-term HSC functionality is a critical barrier in HSC applications for treating hematological disorders. Thus, exploring novel molecules for expanding the HSC pool while maintaining HSC function is needed to improve current BM transplantation strategies.

Epithelial Growth Factor Like 7 (EGFL7) is a ∼30 kDa secreted protein required for efficient angiogenesis (7–9), vasculogenesis (10), and neurogenesis (11). Though EGFL7 is expressed at its highest levels during embryogenesis (12, 13), EGFL7 is constitutively expressed at lower levels in the quiescent endothelium of vessels, and expression is increased upon vascular injury and during active angiogenesis in adults (13). Furthermore, we have recently shown that the EGFL7 is expressed in malignant acute myeloid leukemia (AML) blasts (14) and lymphoma cells (15) and antagonizes the canonical NOTCH signaling in AML (16). However, a role for EGFL7 in normal hematopoiesis HSPCs has not been extensively examined.

Predicted gene expression profiling of EGFL7 in various human tissues shows the highest level of EGFL7 expressions in CD34+ hematopoietic cells (17), suggesting a potential role in HSC regulation. EGFL7 is also highly expressed in cells that comprise the bone marrow microenvironment (BMM) (7), and plays an important role in the regulation of acute graft vs host disease by endothelial cells (18), suggesting a role in HSC-niche interactions. To further characterize a functional role for EGFL7 in the regulation of HSC, we utilized a constitutive, germline *Egfl7* knock-out (*Egfl7* KO) mouse model. Remarkably, *Egfl7* KO mice have reduced BM cellularity and HSC cycling that can be rescued by treating these *Egfl7*KO mice with recombinant EGFL7 (rEGFL7) protein. Interestingly, when we treated WT mice with rEGFL7 protein we observed that EGFL7 can expand HSCs *in vivo* without compromising their functionality. Thus, we have identified EGFL7 as a novel regulator of HSC function with potential translational implications.

## METHODS

### Animal usage

12-16 weeks old, previously generated *Egfl7* KO mice (19) were used for all experiments. Age– and sex-matched C57Bl/6J mice purchased from The Jackson Laboratory (Bar Harbor, ME) were used as WT controls. All animals were housed in the animal facility at The Ohio State University or at Huntsman Cancer Institute, University of Utah. All animal studies were conducted according to protocols approved by the Institutional Animal Care and Use Committees of The Ohio State University and Huntsman Cancer Institute at the University of Utah.

### BrdU incorporation assay

Mice were injected with 5′-bromo-2′-deoxyuridine (BrdU) (BD Bioscience) intraperitoneally (i.p) (50 µg/g of mouse body weight) on day 0. BrdU treatment was continued daily from day 0-30 via drinking water (0.8 mg/ml) until mice were euthanized at day 1-, 3-, 7– or 30– post BrdU injection.

### 5-fluorouracil (5-FU) assays

For the 5-FU survival assay, mice were injected with 5-FU i.p at the dose of 150 µg /mouse/ week and observed for their survival. For 5-FU BM phenotyping, mice were injected once with 5-FU i.p at the dose of 150 µg /mouse. Mice were sacrificed at day 3 or day 7 for HSPC immunophenotyping and cell cycle analysis.

### Recombinant EGFL7 treatment

12-16 weeks old C57/Bl6 mice were injected i.p with 10 µg/mouse of recombinant EGFL7 protein (rEGFL7) (Peprotech) daily for 10 days. Age– and sex-matched C57/Bl6 mice injected with equal volume of PBS were used as controls. Mice were sacrificed on day 11 and BM was analyzed.

## RESULTS

### *Egfl7* KO mice have reduced BM cellularity and reduced number of HSPCs

To investigate the role of EGFL7 on normal hematopoiesis, we evaluated the hematopoietic compartment of the *Egfl7^-/-^* genetic knockout (*Egfl7* KO) mice. First, we validated *Egfl7*-deletion in the HSPCs in these mice by both RNA and protein using qRT-PCR and immunofluorescence chemistry (IFC) respectively (Figure S1A-C). After confirming *Egfl7* knock-out in these mice, we analyzed the BM compartment of the *Egfl7* wild-type (WT) and *Egfl7* KO mice, as this is the primary site of hematopoiesis (20). HSPC immunophenotyping by flow cytometry was performed to determine if differences in the frequency of various HSPC populations was observed (Figure S2A). We found no differences in HSPC frequencies within the BM of *Egfl7* KO mice compared to the age-matched WT mice (Figure S2B-D) with no change in their commitment to various hematopoietic lineages (Figure S2E-F).

However, we did find a significant decrease in BM cellularity of *Egfl7* KO mice compared to age– and sex-matched WT mice resulting in decreases in the total number of long-term (LT) and short-term (ST) HSCs in the BM of *Egfl7* KO mice (Figure 1A-D). Similar reductions were found in the numbers of multi-potent progenitor cells (MPPs), hematopoietic progenitor cells type-1 (HPC), and hematopoietic progenitor cells type-2 (HPC2), as well as committed progenitor cells such as megakaryocyte-erythroid progenitors (MEPs), common myeloid progenitors (CMPs), granulocyte-monocyte progenitors (GMP), and common lymphoid progenitor (CLP) (Figure S2G-I). Reductions in HSCs in *Egfl7* KO mice compared to the WT mice were confirmed using whole mount IFC. Sternal sections from WT and *Egfl7* KO mice were stained using fluorochrome tagged lineage (lin), CD41, and CD48 (shown in green), and CD150 (shown in red) antibodies and imaged under a confocal microscope for quantification of HSCs (Lin^-^ CD48^-^ CD41^-^ CD150^+^) (Fig 1E). IFC imaging revealed a similar reduction in HSCs in *Egfl7* KO mice compared to WT mice (Figure 1F) found by flow cytometry and immunophenotyping. These results indicate that at least one role for Egfl7 in normal hematopoiesis may involve regulating LT-HSCs, reducing the overall numbers of the subsequent ST-HSCs and progenitor numbers.

**Figure 1.**
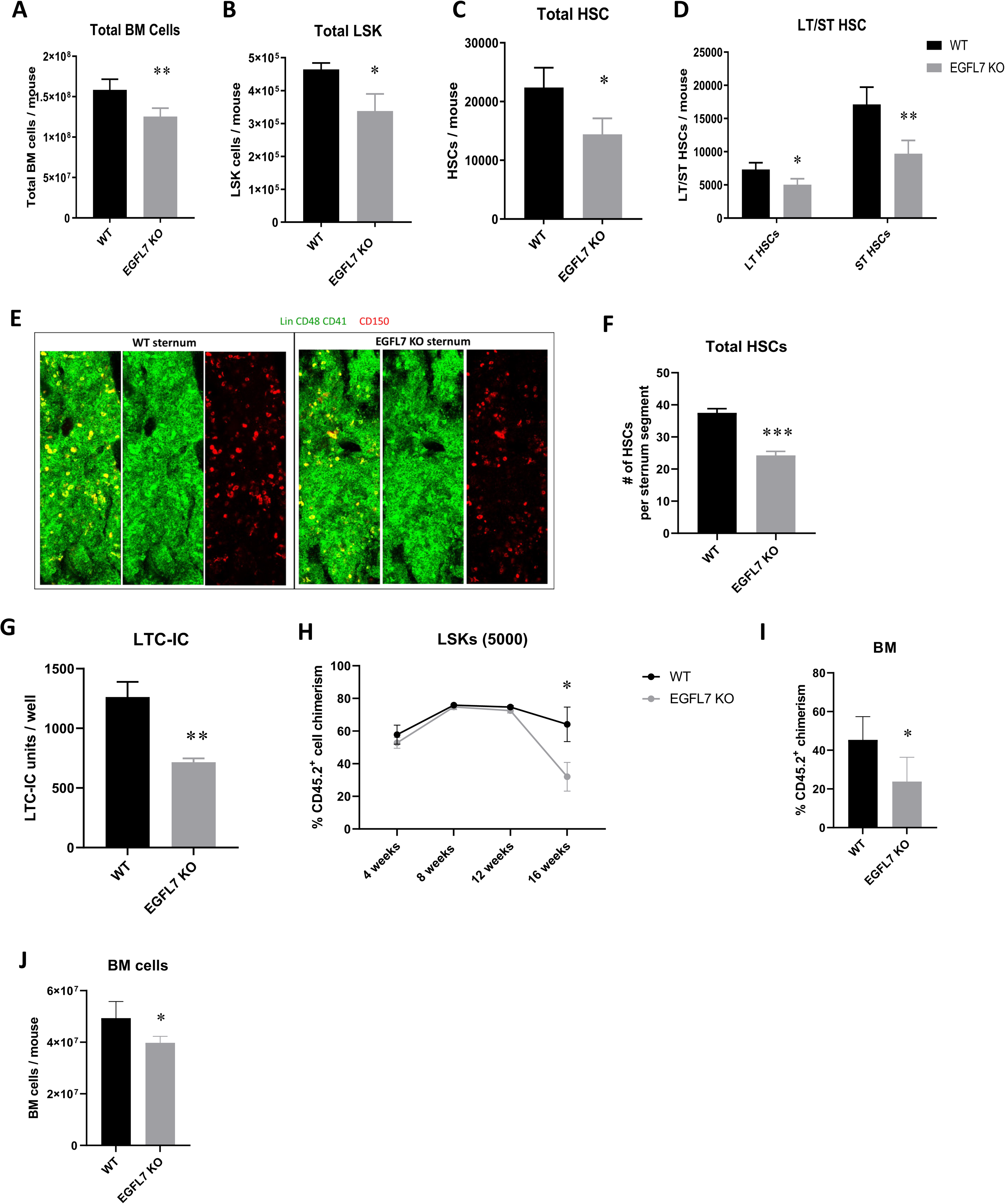
EGFL7 is important for maintaining HSC numbers in the BM. **(A)** 12-16 weeks old WT or *Egfl7* KO mice were sacrificed, and BM cell populations were analyzed by flow cytometry, and LTC-IC assays. Total number of BM cells in *Egfl7* KO mice compared to WT. **(B)** Number of BM LSK (Lin-Sca1+ cKit+ cells), **(C)** number of BM HSCs (Lin-Sca1+ cKit+ CD150+ CD48-cells) and **(D)** number of BM Long-Term (LT) HSCs (Lin-Sca1+ cKit+ CD150+ CD48-CD34-cells) and Short-Term (ST) HSCs (Lin-Sca1+ cKit+ CD150+ CD48-CD34-cells) in *Egfl7* KO mice compared to WT. **(E)** Representative Z-stack immunofluorescence images of sterna from *Egfl7* KO mice stained with Lin (green), CD41 (green), CD48 (green), and CD150 (red) compared to WT. **(F)** Quantification of number of HSCs per sternum segment in *Egfl7* KO mice compared to WT. **(G)** Total number of LTC-IC colonies from BM of *Egfl7* KO mice compared to WT. **(H)** Lethally irradiated WT BoyJ mice were competitively transplanted at 1:1 ratio with sort-purified CD45.2^+^ LSK cells from BM of *Egfl7* KO or WT mice and sort-purified WT CD45.1^+^ LSK cells. Mice were bled every 4 weeks and peripheral blood chimerism was analyzed by flow cytometry. At 16 weeks post transplantation, mice were sacrificed, and BM cells were analyzed by flow cytometry. Percentage of CD45.2^+^ cells in peripheral blood of transplanted mice at 4-16 weeks post transplantation. **(I)** Percentage of CD45.2^+^ BM cells and **(J)** total number of BM cells in transplanted mice at 16 weeks post-transplantation. These data are representative of 3 or more experiments done with N= 4-6 mice per group. *P* = *<0.05, **<0.01, ***<0.001.

### *Egfl*7 KO HSCs have decreased engraftment capacity

To evaluate differences in the functional capacity of *Egfl7* KO and WT HSCs, we performed long term colony-initiating cell (LTC-IC) assays. BM cells from *Egfl7* KO or WT mice were co-cultured on irradiated BM stromal cells for 4 weeks and colony forming cells (CFCs) were performed to estimate the number of HSCs (21). Consistent with our immunophenotyping data, *Egfl7* KO mice had significantly reduced numbers of functional HSCs in their BM compartment (Figure 1G) compared to WT controls. These data support our finding that *Egfl7* loss results in reduced HSC function leading to reduced HSC numbers and overall BM cellularity.

To further examine the role of Egfl7 in HSC function *in vivo*, we performed competitive repopulation unit (CRU) assays. *Egfl7* KO or WT FACS sorted LSK cells (CD45.2^+^) were co-transplanted with equal number of WT competitor LSK cells (CD45.1^+^) into lethally irradiated WT (CD45.1^+^) recipient mice. Analysis of peripheral blood (PB) from recipients at 16 weeks post-transplant, showed a significant decrease in donor-derived chimerism (Figure 1H) as well as the 16-week BM analysis (Figure 1I). BM analysis further revealed that the recipients who received CD45.2^+^*Egfl7* KO LSK cells have significantly less BM cellularity as compared to WT recipients (Figure 1J).

Due to our observation of compromised HSCs in Egfl7 KO mice, we also performed CRU using whole bone marrow (WBM) in which WT (CD45.2+) or *Egfl7* KO (CD45.2+) WBM cells mixed were co-transplanted with WT (CD45.1+) at a ratio of 10:1 (CD45.2+:CD45.1+) into lethally irradiated recipient (CD45.1+) mice. No differences were observed in donor chimerism in the PB 4-16 weeks post-BM transplant (BMT) and within the BM at 16-weeks (Figure 2A-B), We also did not observe any differences in donor chimerism of LSK and HSCs in BM at 16 weeks (Figure 2C-D).

**Figure 2.**
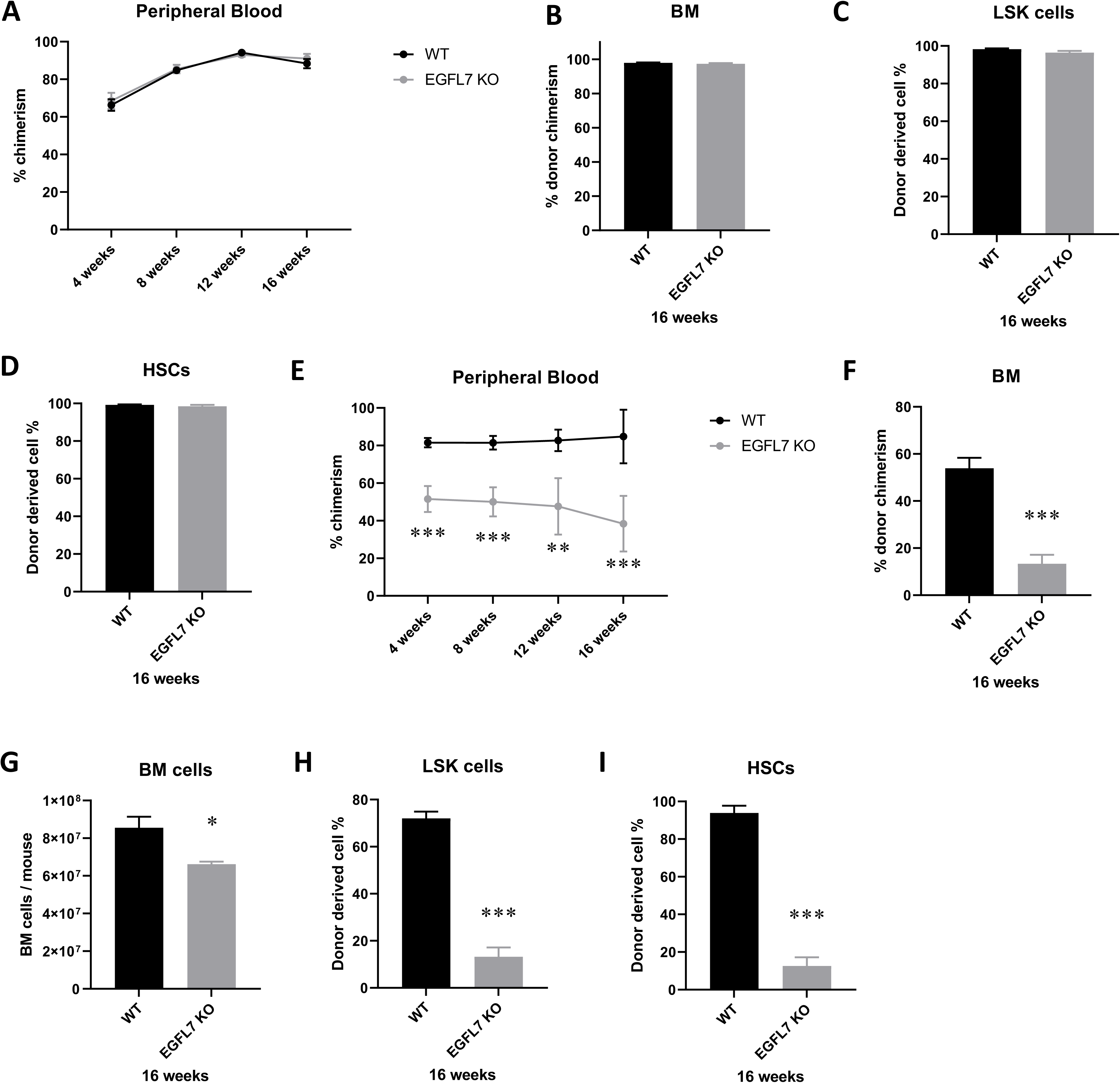
HSCs from *Egfl7* KO mice have reduced serial transplantation capacity. **(A)** Lethally irradiated WT BoyJ mice were transplanted with BM from 12-16 weeks old WT or *Egfl7* KO mice mixed with BM from WT BoyJ mice at 10:1 ratio. Mice were bled every 4 weeks and peripheral blood chimerism was analyzed by flow cytometry. At 16 weeks post transplantation, mice were sacrificed, and BM cells were analyzed by flow cytometry. Percentage of CD45.2^+^ cells in peripheral blood of transplanted mice at 4-16 weeks post-transplantation. **(B)** Percentage of BM CD45.2^+^ cells, **(C)** percentage of CD45.2^+^ BM LSK cells, and **(D)** percentage of CD45.2^+^ BM HSCs in transplanted mice at 16 weeks post-transplantation. **(E)** BM from above primary transplanted mice were transplanted into lethally irradiated WT BoyJ mice. Recipient secondary transplanted mice were bled every 4 weeks and peripheral blood chimerism was analyzed by flow cytometry. At 16 weeks post transplantation, secondary transplanted mice were sacrificed, and BM cells were analyzed by flow cytometry. Percentage of CD45.2^+^ cells in peripheral blood of secondary transplanted mice. **(F)** Percentage of BM CD45.2^+^ cells, **(G)** total number of BM cells, **(H)** percentage of CD45.2^+^ BM LSK cells, and **(I)** percentage of CD45.2^+^ BM HSCs in secondary transplanted mice. These data are representative of 3 or more experiments done with N= 3-6 mice per group. *P* = *<0.05, **<0.01, ***<0.001.

However, to assess whether Egfl7 plays a role in the more primitive HSCs, primary-BMT recipient BM cells were harvested and used to perform secondary-BMT assays. Donor chimerism was assessed in the PB 4-16 weeks post-BMT and within the BM at 16-weeks and demonstrated reduced donor cell chimerism in recipients transplanted with *Egfl7* KO cells (Figure 2E-F). We also found a reduction in overall BM cellularity resulting from decreased percentages of LSK and HSCs in recipients transplanted with *Egfl7* KO (CD45.2^+^) cells (Figure 2G-I). Overall, these results support our findings that loss of *Egfl7* results in impaired serial transplantation capacity and reconstitution potential.

### *Egfl7* loss delays cell cycle progression in HSCs

Our immunophenotypic and transplant data suggests that Egfl7 plays a role in HSCs and might be important for cell cycle regulation and expansion of HSCs. Therefore, we investigated whether *Egfl7* KO HSCs had defects in cell cycle regulation using BrdU incorporation. BrdU was injected into mice (day 0) and added to their drinking water. Mice were analyzed at days: 1, 3, and 7 for BrdU-incorporation in BM populations. We found a significant decrease in proliferating (BrdU+) HSCs, and LSK cells from *Egfl7* KO mice as early as day 3 of BrdU treatment (Figure 3A-C and Figure S3A-C) in *Egfl7* KO mice. To assess whether loss of *Egfl7* results in reduced cycling of LT-HSCs which divide less frequently than ST-HSCs and progenitor cells, mice were dosed for 30 days with BrdU. *Egfl7* KO HSCs had reduced percentage of HSCs in S-phase with a concomitant increase in the number of HSCs found in the G_0_-G_1_ phase compared to WT (Figure 3D). These data suggest that Egfl7 regulates HSC exit from quiescence into cycle in a subset of HSCs. To further validate these findings, WT HSCs were sort-purified and co-stained for EGFL7 and Ki67+ and analyzed by IFC, as previously described (22). We found that the EGFL7 protein levels corresponded with Ki67 expression (Figure 3E-F). Furthermore, WT and *Egfl7* KO HSCs were sort-purified and Ki67 expression levels were determined. The percentage of Ki67+ cells in *Egfl7* KO HSCs were significantly lower compared to WT HSCs (Figure 3G).

**Figure 3.**
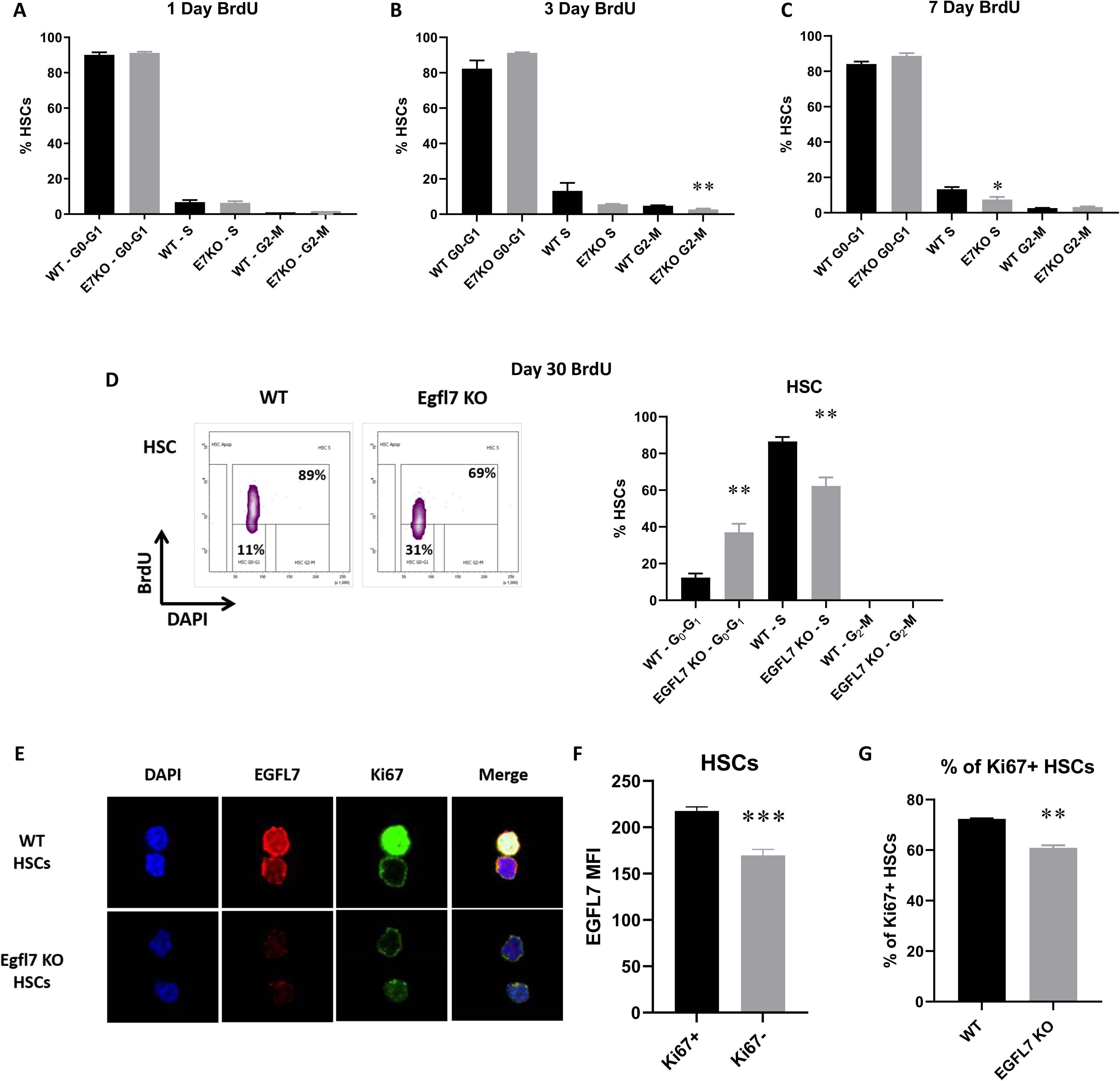
*Egfl7* loss delays cell cycle progression in HSCs. 12-16 weeks old WT or *Egfl7* KO mice were injected with BrdU (50µg/gm, i.p), followed by daily BrdU treatment through drinking water (0.8 mg/ml) until mice were sacrificed at 1-, 3-, 7-, and 30-days post BrdU injection. BM populations were analyzed for BrdU incorporation by flow cytometry. **(A)** Percentage of HSCs in the G_0_-G_1_, S, and G_2_-M cell cycle phases in *Egfl7* KO mice compared to control on day 1, **(B)** 3, and **(C)** 7 post BrdU treatment. Mice were sacrificed 30 days post BrdU treatment and BM was analyzed for BrdU incorporation by flow cytometry. **(D)** (left) Representative flow plots of BrdU incorporation assay at 30 days in BM HSCs from *Egfl7* KO mice compared to WT. (right) Percentage of cells in G_0_-G_1_, S, and G_2_M phases in *Egfl7* KO mice compared to WT. **(E)** BM HSCs were sorted from 12-16 weeks old WT or *Egfl7* KO mice and analyzed for EGFL7 expression by immunofluorescence. Representative immunofluorescence images of BM HSCs stained for EGFL7 (red) and Ki67 (green). Nuclei are counterstained with DAPI. **(F)** Mean fluorescence intensity of EGFL7 in Ki67+ BM HSCs compared to Ki67-HSCs from WT mice. **(G)** Quantification of percentage of Ki67+ BM HSCs in *Egfl7* KO mice compared to WT. These data are representative of 3 or more experiments with N=4-7 mice per group. *P* = *<0.05, **<0.01, ***<0.001.

Taken together, these data show that EGFL7 has a role in the regulation of proliferation of a subset of HSCs and that loss of *Egfl7* results in reduced numbers of cycling HSCs and decreases in BM cellularity without lineage skewing.

### Exogenous EGFL7 treatment increases HSC frequency and enhances engraftment potential

Knowing the important role for EGFL7 in regulating HSC proliferation, we hypothesized that exogenous EGFL7 could increase HSC numbers. To test this, we treated WT mice with recombinant EGFL7 protein (rEGFL7) (Figure 4A). We found BM cellularity and frequency of LSK cells were increased significantly in the rEGFL7 (rE7) treated group compared to vehicle control (CNT) resulting in increased numbers of LT-HSCs, HSCs, and LSKs (Figure 4B-F). However, the frequency of HSCs and LT-HSCs within the LSK population was unchanged (Figure 4G-H) suggesting again that rEGFL7 treatment increases HSCs early in hematopoiesis without downstream perturbations.

**Figure 4.**
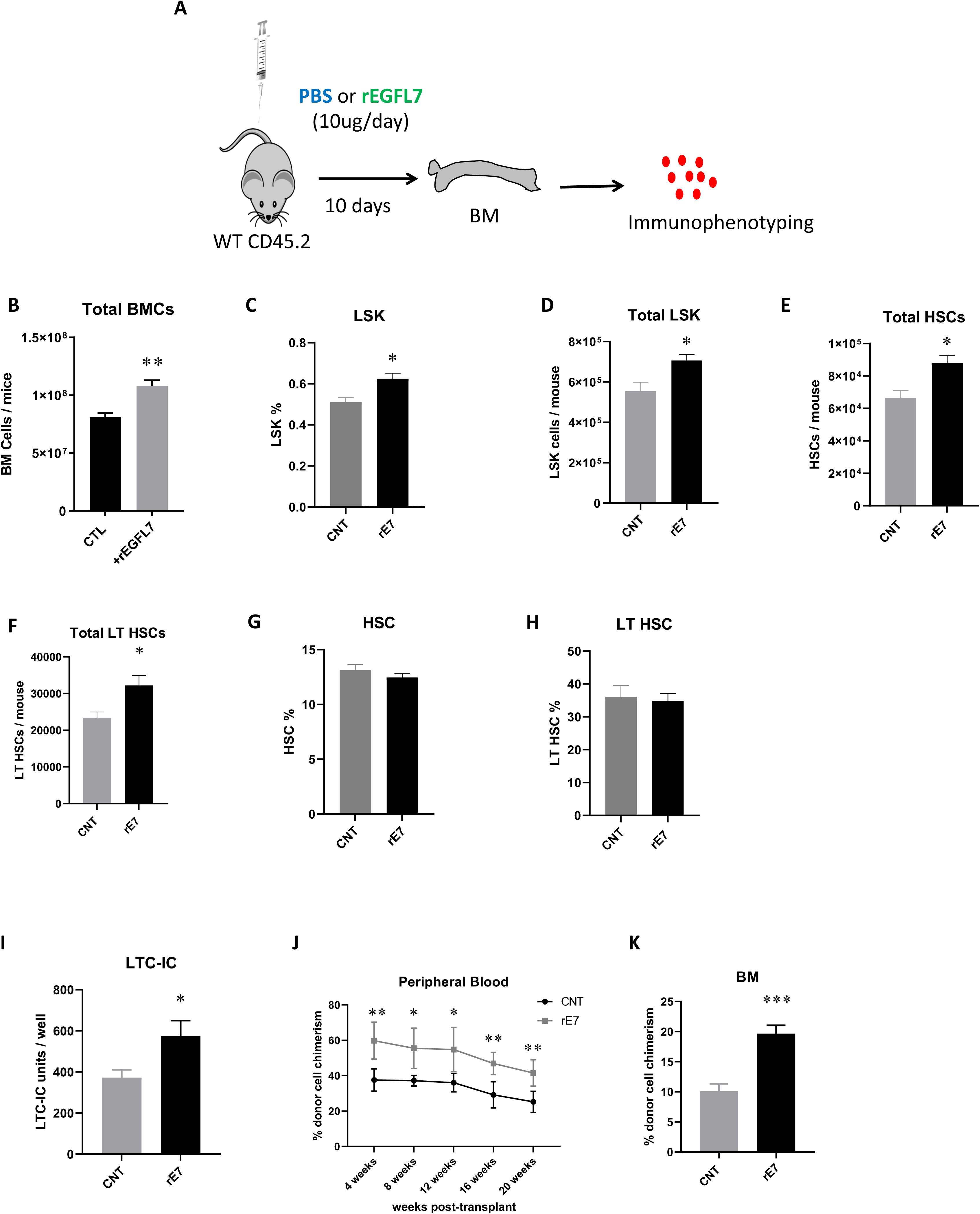
Exogenous EGFL7 treatment increases HSC frequency and enhances engraftment potential. **(A)** Schematic representation of experiment. WT mice were treated with rEGFL7 or PBS (control) for 10 days after which mice were sacrificed, and BM was analyzed by flow cytometry, and LTC-IC assays. **(B)** Total BM cells, **(C)** percentage of BM LSKs, **(D)** total BM LSKs, **(E)** total HSCs, **(F)** total BM LT-HSCs, **(G)** percentage of BM HSCs, and **(H)** percentage of BM LT-HSCs in WT mice 10 days after treatment with rEGFL7 compared to control. **(I)** Total number of LTC-IC colonies in BM from WT mice after treatment with rEGFL7 compared to control for 10 days. **(J)** BM from rEGFL7-treated or control mice was transplanted into lethally irradiated WT BoyJ mice. Mice were bled every 4 weeks and BM was analyzed at 20 weeks post-transplantation. Percentage of CD45.2^+^ cells in peripheral blood of transplanted mice at 4-20 weeks post-transplantation. **(K)** Percentage of CD45.2^+^ cells in BM of transplanted mice at 20 weeks post-transplantation. These data are representative of 3 or more experiments performed in triplicates with N= 3-6 mice per group. *P* = *<0.05, **<0.01, ***<0.001.

Next, we examined whether the increased number of immunophenotypic HSCs *in vivo* results in increased numbers of functional HSCs *in vitro* and *in vivo*. First, we performed LTC-IC assays and found significant increases in the number of LTC-ICs (Figure 4I), representative of functional of HSCs, in mice treated with rEGFL7 compared to CNT. Next, we performed CRU assays to determine if increased numbers of HSCs in the BM leads to increased engraftment after BMT. 2X10^6^ WBM cells from rEGFL7 treated or CNT mice were co-transplanted with 2X10^6^ CD45.1^+^ WBM cells into lethally irradiated WT (CD45.1+) mice. The mice which received rEGFL7-treated donor cells showed a significantly higher percentage of donor cells in their PB and BM (Figure 4J-K). These data suggest that rEGFL7-expanded HSCs maintained their transplantation potential and that the short-term treatment can increase the number of functional HSCs while maintaining preserving HSC functionality.

### Exogenous EGFL7 restores HSC numbers in *Egfl7* KO

Knowing that rEGFL7 treatment resulted in increased HSCs, we next wanted to determine whether the HSC defect observed in the *Egfl7* KO mice could be reversed and rescued with rEGFL7 treatment. We found that rEGFL7 treatment significantly increased BM cellularity of *Egfl7* KO mice. rEGFL7 treatment also increased the LSK population in *Egfl7* KO mice comparable to WT levels (Figure 5A-B). Total HSC numbers in the BM of *Egfl7* KO mice increased significantly after rEGFL7 treatment (Figure 5C-D). Next, BrdU incorporation assay was performed on rEGFL7 treated *Egfl7* KO mice to study if rEGFL7 induces the cycling of *Egfl7* KO HSCs. Indeed, rEGFL7 treatment induces *Egfl7* KO LSK cells to exit G0-G1 phase of cell cycle and enter the cycling state (S phase) (Figure 5E). Overall, our data suggest that EGFL7 is a potent activator of quiescent HSCs.

**Figure 5.**
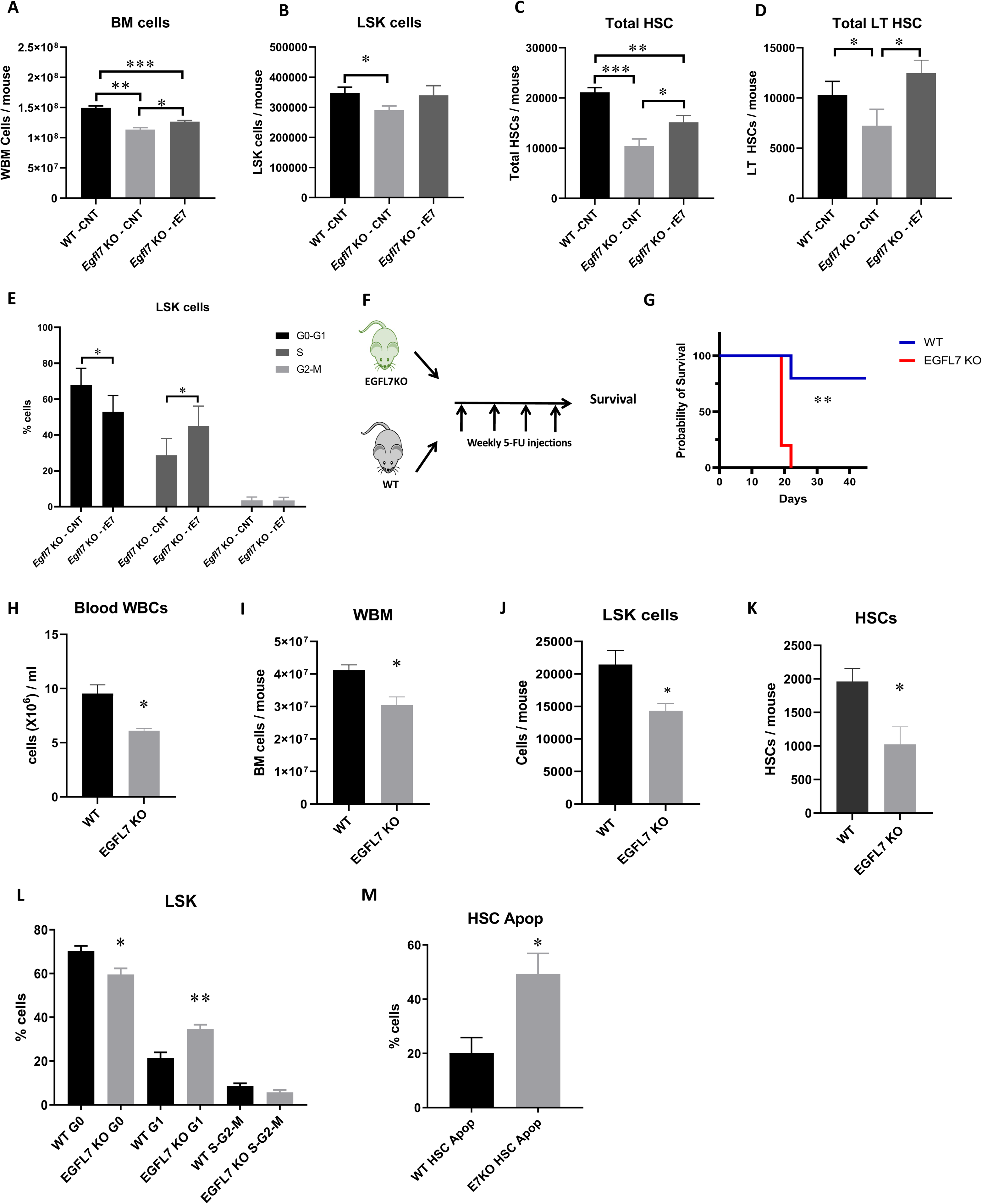
Exogenous EGFL7 restores HSC numbers in *Egfl7* KO and protects HSCs in irradiation stress. **(A)** 12-16 weeks old *Egfl7* KO mice were treated with rEGFL7 (Egfl7 KO – rE7) or PBS (Egfl7 KO – CNT) for 10 days after which BM was analyzed by flow cytometry. WT mice treated with PBS were used as controls (WT – CNT). Total number of BM cells, **(B)** total number of BM LSKs, **(C)** total number of BM HSCs, **(D)** total number of BM LT-HSCs in Egfl7 KO – rE7 and Egfl7 KO – CNT mice compared to WT – CNT. **(E)** 12-16 weeks old *Egfl7* KO mice were treated with rEGFL7 (Egfl7 KO – rE7) or PBS (Egfl7 KO – CNT) for 10 days. For the BrdU assay, mice were injected with BrdU (50µg/gm, i.p) on day 0, followed by daily BrdU treatment through drinking water (0.8 mg/ml) until mice were sacrificed on day 11 and BM was analyzed for BrdU incorporation by flow cytometry. Percentage of BM LSK cells in G_0_-G_1_, S, and G_2_M phases in in Egfl7 KO – rE7 and Egfl7 KO – CNT mice compared to WT – CNT. **(F)** Schematic representation of 5-FU survival assay. 12-16 weeks old WT or *Egfl7* KO mice were injected with 5-FU (150 mg/kg, i.p.) once a week for 4 weeks. Mice were monitored daily for survival. **(G)** Kaplan-Meier survival curve of WT and *Egfl7* KO mice after 5-FU treatment. **(H)** 12-16 weeks old WT or *Egfl7* KO mice injected with 5-FU (150 mg/kg, i.p.) were sacrificed 3 days post 5-FU injection and peripheral blood and BM was analyzed by an automated hematology analyzer. BM cells were stained with Ki67 and DAPI to analyze proliferating LSK cells, or with Annexin V to analyze apoptotic cells by flow cytometry. Number of peripheral blood WBCs in peripheral blood, **(I)** number of BM cells, **(J)** number of BM LSK cells, **(K)** number of BM HSCs in WT or *Egfl7* KO mice on day 3 after 5-FU treatment. **(L)** Percentage of BM LSK cells in G_0_-G_1_, S, and G_2_M phases in WT or *Egfl7* KO mice. **(M)** Percentage of apoptotic BM HSCs in WT or *Egfl7* KO mice. These data represent 3 or more experiments performed in triplicates with N= 3-6 mice per group. *P* = *<0.05, **<0.01, ***<0.001.

### Egfl7 protects HSCs in 5-FU and irradiation stress

Recognizing the importance of EGFL7 in recruiting the HSCs to active cell cycle, we challenged the *Egfl7* KO mice with 5-fluorouracil (5-FU) to induce hematopoietic stress. As HSCs are stimulated to cycle rapidly after 5-FU treatment (5), we injected *Egfl7* KO mice with a weekly dose of 5-FU (150 mg/kg). These mice were followed for their survival for 4 weeks continuously (Figure 5F). *Egfl7* KO mice failed to survive after the third 5-FU injection, while most of the WT mice survived the weekly 5-FU treatment, indicating that the *Egfl7* KO mice were highly susceptible to induced hematopoietic stress (Figure 5G). Moreover, WT HSCs are stimulated to proliferate rapidly within 3 days and then stabilize at 7 days post-treatment (5). We analyzed the BM HSC compartment of *Egfl7* KO mice at 3 days and 7 days post-5-FU injection (150mg/kg). Peripheral white blood counts (WBC) were lower in *Egfl7* KO mice compared to WT counterparts at 3 days post-5-FU treatment (Figure 5H), confirming a delay in replenishing mature hematopoietic cells in *Egfl7* KO mice. Analysis of the BM compartment of these mice on day 3 showed reduced BM cellularity (Figure 5I). The number of LSK cells and HSCs were also considerably reduced in these mice (Figure 5J-K). The BM cells from 5-FU treated *Egfl7* KO mice, when analyzed for their cell cycle status using Ki67/DAPI, showed an increased frequency of LSK cells in the G1 phase of the cell cycle (Figure 5L). We also analyzed apoptotic HSCs, as previously described (23), and found that percentage of apoptotic HSCs in *Egfl7* KO BM was significantly increased at day 3 post-5-FU treatment as compared to 5-FU treatment WT mice BM HSCs (Figure 5M). Similar analysis of BM of *Egfl7* KO mice at day 7 post-5-FU treatment shows a significantly low number of total BM cells along with a reduced number of LSK cells, HSCs, LT, and ST-HSCs (Figure S3D-H). These data taken together show that the EGFL7 is required for the proliferation of HSCs and for restoration of homeostasis after 5-FU induced stress conditions.

We also assessed whether rEGFL7 could facilitate the recovery of HSCs after irradiation induced hematopoietic stress. For these experiments, WT mice were given a sublethal dose of irradiation (5Gy). Mice were then treated with either rEGFL7 (10µg/day) or vehicle control (CNT) for 10 days. Although rEGFL7 treatment did not alter the BM cell count (Figure S3I) or percentage LSK cells (Figure S3J), the total number LSK cells were significantly higher in rEGFL7 treated mice (Figure S3K). MPP, HPC1 and HPC2 cell numbers were also seen to be enhanced significantly in rEGFL7 treated mice after irradiation (Figure S3L). These data suggest that EGFL7 plays a role in HSC recovery after genotoxic stresses such as 5-FU and irradiation, further supporting its importance in regulating HSC numbers, functionality, transplantation, and protection during genotoxic stress.

### scRNAseq analysis reveals dysregulation of cell cycle regulation in response to altered EGFL7 activity

To better understand the molecular pathways regulated by Egfl7 in HSCs, we performed scRNAseq on HSPC cells (cKit^+^ selected cells) isolated from the BM of *Egfl7* KO and WT mice (Figure S4A). Unbiased clustering of cKit^+^ cell expression profiles resulted in 20 different clusters of hematopoietic cells (Figure 6A, Figure S4B). Key marker gene expression was used to annotate each cluster type (24). Cluster 3 was annotated as HSC due to the high expression of the HSC-specific gene *Hlf*. Similarly, multiple clusters were annotated as MPP2, MPP3, and MPP4 based on high expression of *Pf4*, *Ctsg,* and *Dntt* respectively (24) (Fig 6B). Mature hematopoietic cell populations were identified using a reference map for hematopoiesis (24). The following clusters were annotated based on the high expression of these mature cell-specific marker genes: *Fcrla* / *Vpreb1* (B cells), *Cd74* (Dendritic cells), *Ccl5* (Innate lymphocytes), *Siglech* (Plasmocytes), *Ly86* (Monocytes), *Fcnb* / *Ltf* / *Gstm1* (Neutrophils), *C1qb* (Macrophages), *Prg2* (Eosinophils), *Prss34* (Basophils) and *Hba-a2* / *Car1* (Erythrocytes) (Figure S4C).

**Figure 6.**
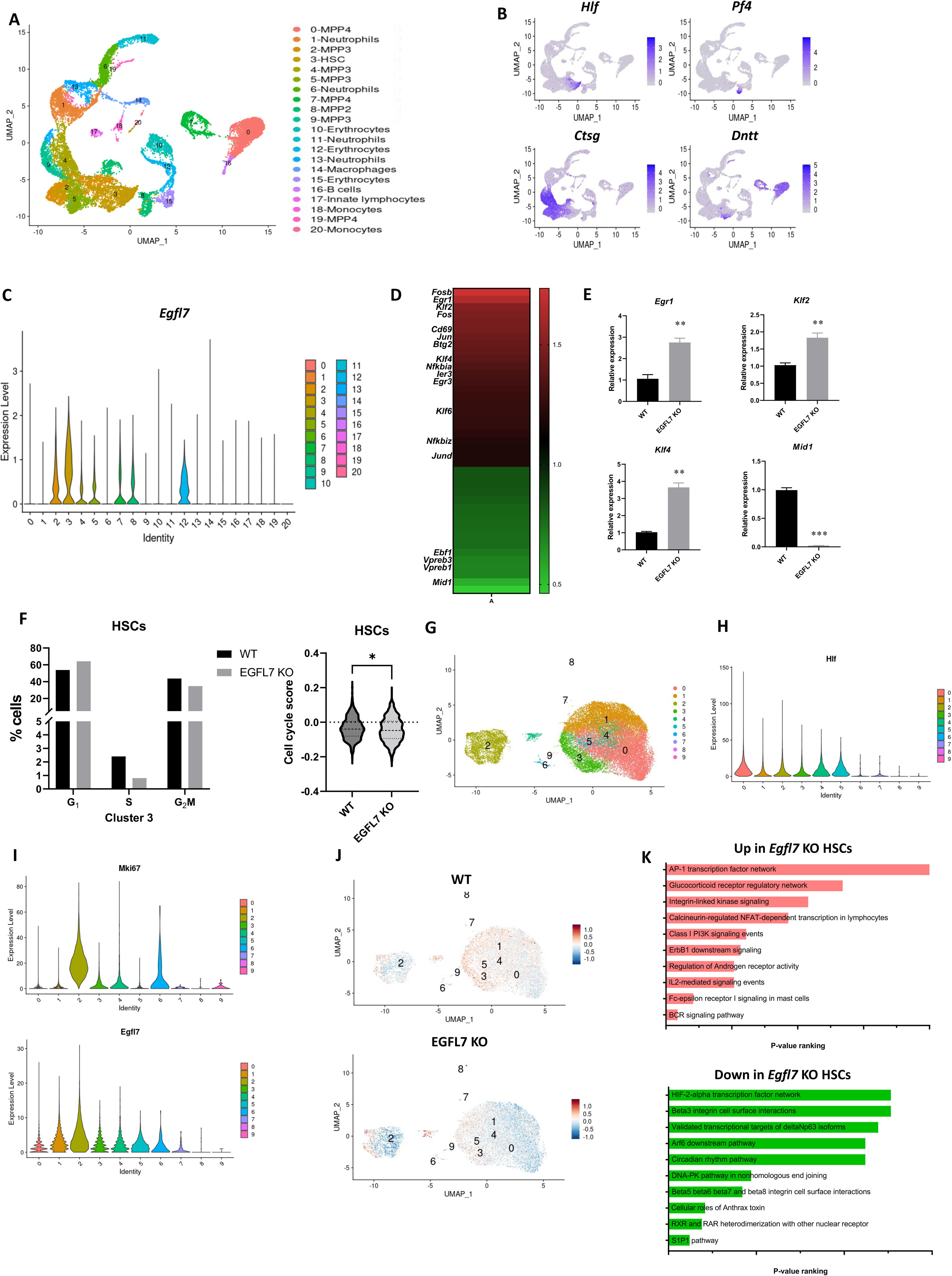
scRNAseq analysis reveals dysregulation of cell cycle regulation in response to altered EGFL7 activity. **(A)** BM cKit^+^ cells (HSPCs) from 12-16 weeks old WT and *Egfl7* KO mice were subjected to scRNA-sequencing. Uniform Manifold Approximation and Projection (UMAP) of HSPCs from BM of *Egfl7* KO and WT mice, colored by clustering and annotated based on gene signature. **(B)** UMAP of HSPCs colored by expression of key cell-type marker genes *Hlf* (HSCs), *Pf4* (MPP2), *Ctsg* (MPP3), and *Dntt* (MPP4). **(C)** Violin plot distribution of *Egfl7* expression levels across all HSPC clusters. **(D)** Heatmap of differentially expressed genes in HSC cluster from *Egfl7* KO mice compared to WT. **(E)** Relative mRNA expression of *Egr1*, *Klf2*, *Klf4*, and *Mid1* in HSCs sorted from *Egfl7* KO mice compared to WT, as determined by qRT-PCR. mRNA expression is calculated relative to *Gapdh*. N= 5 mice per group. **(F)** (left) Percentage of HSCs (cluster 3) in G1, S, and G2M cell cycle phases in *Egfl7* KO mice compared to WT. (right) Violin plots of cell cycle scores in HSCs (cluster 3) from *Egfl7* KO mice compared to WT. **(G)** UMAP of BM HSCs (LSK CD150+ cells) from *Egfl7* KO and WT mice, colored by clustering. **(H)** Violin plot distribution of *Hlf,* **(I)** *Mki67* and *Egfl7* expression levels across all HSC clusters. **(J)** UMAP of HSCs colored by cell cycle scores in WT (top) and *Egfl7* KO (bottom) mice. **(K)** Top differentially up– and down-regulated signaling pathways in *Egfl7* KO HSCs compared to WT. *P* = *<0.05, **<0.01, ***<0.001.

We analyzed the expression of *Egfl7* in the unbiased clustered data set. Overlaying the *Egfl7* expression onto the clustered data set we found that the expression was highest in our annotated HSC cluster (Figure 6C). Unbiased sub-clustering of our annotated HSC cluster showed no differential patterns. Subsequent differential gene expression analysis of the HSC cluster revealed that *Egfl7* KO HSCs overexpress genes that regulate quiescence (*Egr1*, *Btg2*, *Egr3*, *Ier3*) (25, 26), self-renewal (*Klf2*, *Klf4*, *Klf6*) (27) and apoptotic markers-AP-1 complex molecules (*Fosb*, *Fos*, *Jun*, *Jund*) (28), along with markers for aged HSCs (*Cd69*, *Nfkbia,* and *Nfkbiz*) (24). *Egfl7* KO HSCs also showed downregulation of B cell lineage commitment marker expression (*Mid1*, *Vpreb1,* and *Vpreb3*) (Figure 6D) (29). Additionally, we confirmed by qPCR the upregulation of *Egr1*, *Klf2,* and *Klf4* and the downregulation of *Mid1,* in sort purified *Egfl7* KO HSCs as compared to WT HSCs (Figure 6E).

Since our data suggests a role for Egfl7 in regulating proliferation/quiescence and self-renewal of HSCs, we also analyzed the cell cycle status in the WT and *Egfl7* KO BM cells using their gene expression profiles (using Seurat cell-cycle scoring and regression workflow). Corresponding to our *in vitro* and *in vivo* data, higher percentages of *Egfl7* KO HSCs (cluster 3) were found to be in G_1_ phase of the cell cycle and a corresponding decreased frequency of the cells in the S-G_2_-M phase was observed. A comparison of the cell cycle score from WT and *Egfl7* KO HSC clusters also indicates decreased cycling of *Egfl7* KO HSCs as compared to their WT counterparts (Figure 6F).

Next, to gain deeper insights into transcriptional changes specific to HSCs, we sort purified LSK CD150+ HSCs from WT and *Egfl7* KO mice and performed scRNA-seq on these cells. Unbiased clustering of WT and *Egfl7* KO LSK CD150+ cells resulted in 9 different clusters (Figure 6G). Cluster numbers 0, 1, 2, 3, 4 and 5, which made up more than 99% of the analyzed cells, expressed significant levels of *Hlf*, confirming their HSC identity (Figure 6H). Interestingly, when the HSC subclusters were analyzed for *Mki67* expression, HSC cluster 2 which showed highest level of *Mki67* expression, also had the highest level of *Egfl7* expression (Figure 6I). In this analysis, WT and *Egfl7* KO HSC clusters were also analyzed for their cell cycle scores. *Egfl7* KO cells demonstrated a clear reduction in their cell cycle score again indicating defects in their cycling status (Figure 6J). Pathway enrichment analysis using the differentially expressed genes from *Egfl7* KO vs WT LSK CD150+ cells show significant induction of AP-1 and reduced HIF-2-alpha and HIF-1-alpha transcription networks (Figure 6K). Analysis of *EGFL7* expression in human hematopoietic cells reveals it is highly expressed in the cycling CD34^+^ HSCs, hinting at its role in regulation of cell cycle in HSCs (30).

### Profiling the rEGFL7 induced transcriptional changes in HSCs

To understand the transcriptional changes associated with rEGFL7 treatment on HSCs, we profiled cKit^+^ cells from WT mice treated with vehicle control (CNT) or rEGFL7 (rE7) for 10 days by scRNA-seq (Figure S5A). Unbiased UMAP clustering of cKit^+^ cells resulted in 23 different clusters (Figure 7A). Unbiased cluster representation showed no evident differences in the CNT (black) and rEGFL7 (red) groups (Figure 7B). We further teased out the top 50 differentially expressed genes among the 23 clusters (Figure S5B). Annotation of the clusters was done based on the specific expression marker for each cell type as described earlier in the study. Representative key cell typing genes are shown for each of the main cluster types: HSC (*Hlf*), MPP2 (*Pf4*), MPP3 (*Ctsg*), and MPP4 (*Dntt*) (Figure 7C). Violin plot for *Hlf* expression further highlights the HSC cluster (Figure 7D). Differentiated and committed cell types were annotated using the same reference map described earlier (Figure S5C).

**Figure 7.**
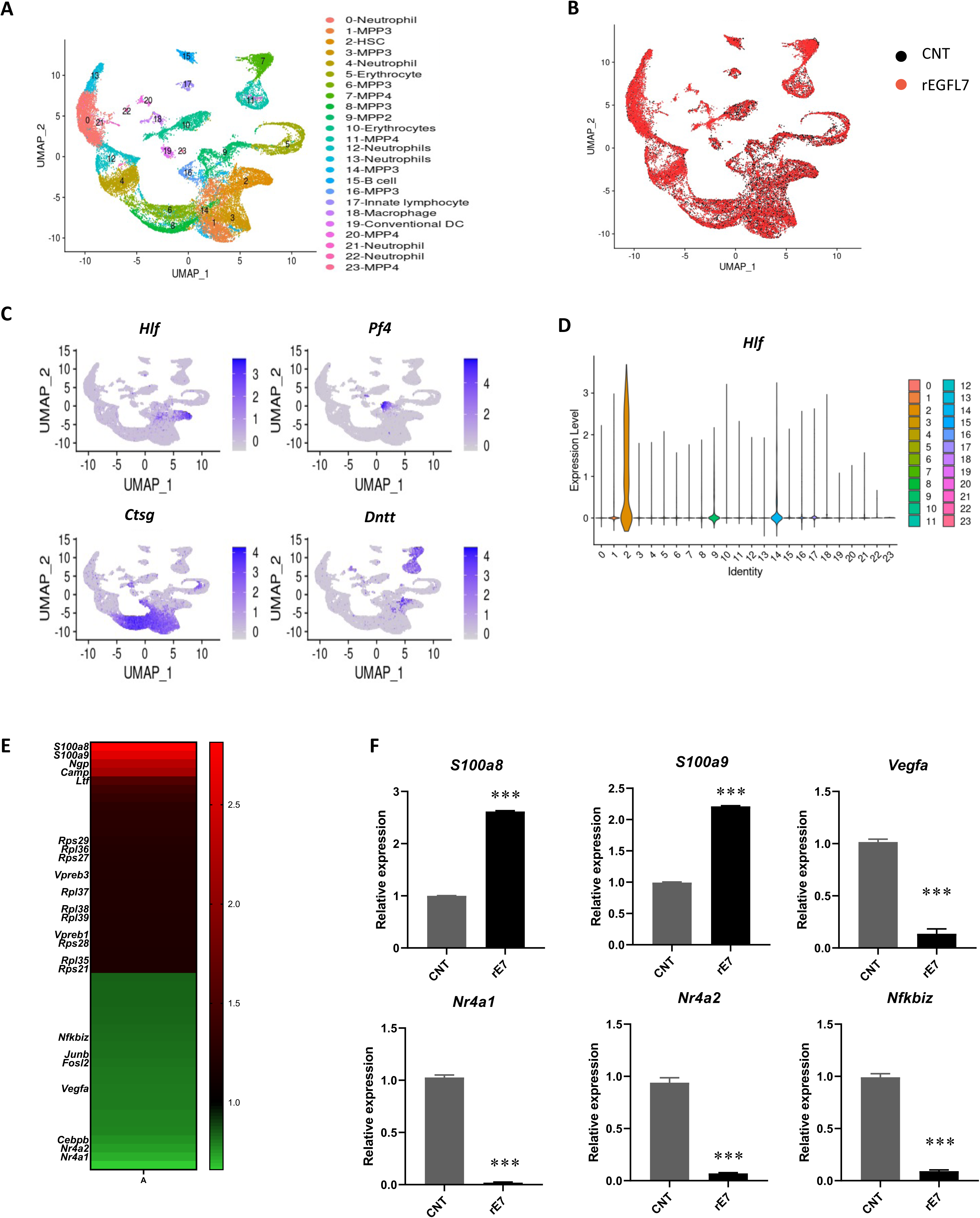
Profiling the rEGFL7 induced transcriptional changes in HSCs. (A) WT mice were treated with rEGFL7 or PBS (control) for 10 days after which BM cKit+ cells (HSPCs) were subjected to scRNA-seq. UMAP of HSPCs from BM of rEGFL7-treated and control mice, colored by clustering and annotated based on gene signature. (B) UMAP of HSPCs colored by treatment group-rEGFL7-treated (red) and control (black). **(C)** UMAP of HSPCs colored by expression of key cell-type marker genes *Hlf* (HSCs), *Pf4* (MPP2), *Ctsg* (MPP3), and *Dntt* (MPP4). **(D)** Violin plot distribution of *Hlf* expression levels across all HSPC clusters. **(E)** Heatmap of differentially expressed genes in HSC cluster from rEGFL7-treated mice compared to control. **(F)** Relative mRNA expression of *S100a8*, *S100a9*, *Vegfa*, *Nr4a1*, *Nr4a2*, and *Nfkbiz* in HSCs sorted from rEGFL7-treated mice compared to control, as determined by qRT-PCR. mRNA expression is calculated relative to *Gapdh*. N=5 mice per group. *P* = *<0.05, **<0.01, ***<0.001.

Analysis for changes in intrinsic gene expression programs showed HSCs from rEGFL7 treated mice had higher expression of proinflammatory molecules such as *S100a8*, *S100a9* (31, 32), *Camp* (33), *Ngp* (34), and *Ltf* (35), indicating that the rEGFL7 treatment induces proinflammatory-like response in the HSC transcriptome (36). HSCs from rEGFL7 treated mice also show increased expression of ribosomal proteins (*Rps21, Rps27, Rps28, Rps29, Rpl35, Rpl36, Rpl37, Rpl38*, and *Rpl39*), which are indicators of active biogenesis in the cells and are implicated in the regulation of apoptosis, cell cycle, and proliferation (37). Simultaneous downregulation of *Nr4a1* and *Nr4a2* also suggests that rEGFL7 induces quiescent HSCs to activate proliferation cascades (38). Overall, cells from the rEGFL7 treatment group showed the exact opposite of what was seen in the Egfl7 KO HSCs, showing increases in *Vpreb1* and *Vpreb3* expression, which are associated with initiation of pre-B-cell-like transcription (29). Similarly, these cells had downregulation of aged HSC markers such as *Nfkbiz* and *Junb* (Figure 7E).

We validated the scRNA-seq results in sort purified HSCs from CNT and rEGFL7 treated mice by qRT PCR. As expected, HSCs from rEGFL7 treated mice demonstrated high expression levels of *S100a8* and *S100a9* and reduced expression of *Nr4a1* and *Nr4a2* (Figure 7F).

## DISCUSSION

EGFL7 has previously been shown to play a role in self-renewal and biology of neuronal and embryonic stem cells (39, 40). These observations led us to hypothesize that EGLF7 could also be essential for normal regulation of HSCs. Using an *Egfl7* KO murine model, we report for the first time that EGFL7 is important for maintaining HSC numbers in the adult BM. Although, *Egfl7* KO mice have normal HSC frequencies in homeostatic conditions, we found that these mice have reduced BM cellularity and total HSCs numbers. We validated these defects in HSCs of Egfl7 KO mice both *in vitro* and *in vivo*. scRNA-seq revealed differential gene expression within the HSCs resulting from the loss of Egfl7. *Egfl7* KO HSCs had increased levels of key regulators of HSC homeostasis such as *Egr1*. Upregulation of *Egr1* along with *Fos*, *Jun*, *Junb*, *Klf4*, *Btg2*, and *Nfkbia* has been shown to be dysregulated in HSCs with impaired cell division (25) and corresponds to the transcriptional changes we found in *Egfl7* KO HSCs providing a possible mechanism to account for the increased quiescence of *Egfl7* KO HSCs.

Treatment of WT mice with rEGFL7 significantly increased HSCs and re-transplantation of HSCs from rEGFL7 treated mice resulted in higher chimeric output without overt lineage skewing. These results are important since it suggests that EGFL7, a secreted protein, can be used as a novel growth factor for expanding HSCs, without compromising HSC functional output. scRNA-seq data of rEGFL7 treated HSCs revealed upregulation of ribosomal proteins, indicative of HSC activation and proliferation. Furthermore, the defects in Egfl7 KO mice in response to 5-FU treatment suggests that at least one role for Egfl7 in HSCs is during hematopoietic stress.

Importantly, loss of Egfl7 does not lead to overt BM failure or premature death in adult mice. This along with the ability of rEGFL7 protein to rescue *Egfl7* KO defects, suggest that Egfl7 loss may exert its effects on only a subpopulation of HSCs and not all BM HSCs. This could be due to specific HSC functions, such HSCs involved in stress response, or possibly HSCs located in different regions within the BM niche. Better resolution of the heterogeneity and regulatory pathways within HSCs will allow us to better tailor *in vitro* and *in vivo* conditions which allow HSCs to better retain functionality in different physiological conditions, improving their use in diverse settings including cellular transplantation therapies.

## Acknowledgments

This work was supported by the National Cancer Institute (NCI), National Institutes of Health (NIH), under the award R01CA259182, and by the National Heart, Lung, and Blood Institutes (NHLBI), National Institutes of Health (NIH) under the award R01HL163849.

## Author Contributions

RK and AMD designed experiments; RK, CG, AR, MK, APU, YB, OB, WL, SC, and KAR performed experiments; EARG and KEM contributed new reagents/analytic tools; RK, CG, and KEM analyzed data; RK, CG, and AMD wrote and, CG, AP, PGP, SE, RG and AMD reviewed and edited the manuscript.

## Data Availability

Data files will be made available upon reasonable request to the corresponding author.

## REFERENCES

1. Spangrude GJ, Heimfeld S, Weissman IL. Purification and characterization of mouse hematopoietic stem cells. Science. 1988;241(4861):58–62.

2. Orkin SH, Zon LI. Hematopoiesis: an evolving paradigm for stem cell biology. Cell. 2008;132(4):631–44.

3. Bradford GB, Williams B, Rossi R, Bertoncello I. Quiescence, cycling, and turnover in the primitive hematopoietic stem cell compartment. Exp Hematol. 1997;25(5):445–53.

4. Nakamura-Ishizu A, Takizawa H, Suda T. The analysis, roles and regulation of quiescence in hematopoietic stem cells. Development. 2014;141(24):4656–66.

5. Harrison DE, Lerner CP. Most primitive hematopoietic stem cells are stimulated to cycle rapidly after treatment with 5-fluorouracil. Blood. 1991;78(5):1237–40.

6. Randall TD, Weissman IL. Phenotypic and functional changes induced at the clonal level in hematopoietic stem cells after 5-fluorouracil treatment. Blood. 1997;89(10):3596–606.

7. Chim SM, Kuek V, Chow ST, Lim BS, Tickner J, Zhao J, et al. EGFL7 is expressed in bone microenvironment and promotes angiogenesis via ERK, STAT3, and integrin signaling cascades. J Cell Physiol. 2015;230(1):82–94.

8. Nichol D, Shawber C, Fitch MJ, Bambino K, Sharma A, Kitajewski J, et al. Impaired angiogenesis and altered Notch signaling in mice overexpressing endothelial Egfl7. Blood. 2010;116(26):6133–43.

9. Nikolic I, Plate KH, Schmidt MHH. EGFL7 meets miRNA-126: an angiogenesis alliance. J Angiogenes Res. 2010;2(1):9.

10. Nikolic I, Stankovic ND, Bicker F, Meister J, Braun H, Awwad K, et al. EGFL7 ligates αvβ3 integrin to enhance vessel formation. Blood. 2013;121(15):3041–50.

11. Bicker F, Vasic V, Horta G, Ortega F, Nolte H, Kavyanifar A, et al. Neurovascular EGFL7 regulates adult neurogenesis in the subventricular zone and thereby affects olfactory perception. Nat Commun. 2017;8:15922.

12. Parker LH, Schmidt M, Jin SW, Gray AM, Beis D, Pham T, et al. The endothelial-cell-derived secreted factor Egfl7 regulates vascular tube formation. Nature. 2004;428(6984):754–8.

13. Nichol D, Stuhlmann H. EGFL7: a unique angiogenic signaling factor in vascular development and disease. Blood. 2012;119(6):1345–52.

14. Papaioannou D, Shen C, Nicolet D, McNeil B, Bill M, Karunasiri M, et al. Prognostic and biological significance of the proangiogenic factor EGFL7 in acute myeloid leukemia. Proc Natl Acad Sci U S A. 2017;114(23):E4641–E7.

15. Goda C, Kolovich S, Rudich A, Karunasiri M, Kulkarni R, Rajgolikar G, et al. Epidermal growth factor-like 7 is a novel therapeutic target in mantle cell lymphoma. Exp Hematol. 2023;123:28–33.e3.

16. Bill M, Pathmanathan A, Karunasiri M, Shen C, Burke MH, Ranganathan P, et al. EGFL7 Antagonizes NOTCH Signaling and Represents a Novel Therapeutic Target in Acute Myeloid Leukemia. Clin Cancer Res. 2020;26(3):669–78.

17. Hong G, Kuek V, Shi J, Zhou L, Han X, He W, et al. EGFL7: Master regulator of cancer pathogenesis, angiogenesis and an emerging mediator of bone homeostasis. J Cell Physiol. 2018;233(11):8526–37.

18. Dorrance AM, Moutuou MM, Goda C, Sell NE, Kalyan S, Karunasiri M, et al. Modulating endothelial cells with EGFL7 to diminish aGVHD after allogeneic bone marrow transplantation in mice. Blood Adv. 2022;6(7):2403–8.

19. Kuhnert F, Mancuso MR, Hampton J, Stankunas K, Asano T, Chen CZ, et al. Attribution of vascular phenotypes of the murine Egfl7 locus to the microRNA miR-126. Development. 2008;135(24):3989–93.

20. Gulati GL, Ashton JK, Hyun BH. Structure and function of the bone marrow and hematopoiesis. Hematol Oncol Clin North Am. 1988;2(4):495–511.

21. Dorrance AM, Neviani P, Ferenchak GJ, Huang X, Nicolet D, Maharry KS, et al. Targeting leukemia stem cells in vivo with antagomiR-126 nanoparticles in acute myeloid leukemia. Leukemia. 2015;29(11):2143–53.

22. Goda C, Kulkarni R, Bustos Y, Li W, Rudich A, Balcioglu O, et al. Cellular taxonomy of the preleukemic bone marrow niche of acute myeloid leukemia. Leukemia. 2025;39(1):51–63.

23. Bill M, Goda C, Pepe F, Ozer HG, McNeil B, Zhang X, et al. Targeting BRD4 in acute myeloid leukemia with partial tandem duplication of the. Haematologica. 2021;106(9):2527–32.

24. Giladi A, Paul F, Herzog Y, Lubling Y, Weiner A, Yofe I, et al. Single-cell characterization of haematopoietic progenitors and their trajectories in homeostasis and perturbed haematopoiesis. Nat Cell Biol. 2018;20(7):836–46.

25. Desterke C, Bennaceur-Griscelli A, Turhan AG. EGR1 dysregulation defines an inflammatory and leukemic program in cell trajectory of human-aged hematopoietic stem cells (HSC). Stem Cell Res Ther. 2021;12(1):419.

26. Min IM, Pietramaggiori G, Kim FS, Passegué E, Stevenson KE, Wagers AJ. The transcription factor EGR1 controls both the proliferation and localization of hematopoietic stem cells. Cell Stem Cell. 2008;2(4):380–91.

27. Park CS, Lewis A, Chen T, Lacorazza D. Concise Review: Regulation of Self-Renewal in Normal and Malignant Hematopoietic Stem Cells by Krüppel-Like Factor 4. Stem Cells Transl Med. 2019;8(6):568–74.

28. Oedekoven CA, Belmonte M, Bode D, Hamey FK, Shepherd MS, Che JLC, et al. Hematopoietic stem cells retain functional potential and molecular identity in hibernation cultures. Stem Cell Reports. 2021;16(6):1614–28.

29. Kudo A, Melchers F. A second gene, VpreB in the lambda 5 locus of the mouse, which appears to be selectively expressed in pre-B lymphocytes. EMBO J. 1987;6(8):2267–72.

30. Bagger FO, Sasivarevic D, Sohi SH, Laursen LG, Pundhir S, Sønderby CK, et al. BloodSpot: a database of gene expression profiles and transcriptional programs for healthy and malignant haematopoiesis. Nucleic Acids Res. 2016;44(D1):D917–24.

31. Foell D, Wittkowski H, Vogl T, Roth J. S100 proteins expressed in phagocytes: a novel group of damage-associated molecular pattern molecules. J Leukoc Biol. 2007;81(1):28–37.

32. Gopal R, Monin L, Torres D, Slight S, Mehra S, McKenna KC, et al. S100A8/A9 proteins mediate neutrophilic inflammation and lung pathology during tuberculosis. Am J Respir Crit Care Med. 2013;188(9):1137–46.

33. Hemshekhar M, Piyadasa H, Mostafa D, Chow LNY, Halayko AJ, Mookherjee N. Cathelicidin and Calprotectin Are Disparately Altered in Murine Models of Inflammatory Arthritis and Airway Inflammation. Front Immunol. 2020;11:1932.

34. Cassatella MA, Östberg NK, Tamassia N, Soehnlein O. Biological Roles of Neutrophil-Derived Granule Proteins and Cytokines. Trends Immunol. 2019;40(7):648–64.

35. Baveye S, Elass E, Mazurier J, Spik G, Legrand D. Lactoferrin: a multifunctional glycoprotein involved in the modulation of the inflammatory process. Clin Chem Lab Med. 1999;37(3):281–6.

36. Freire PR, Conneely OM. NR4A1 and NR4A3 restrict HSC proliferation via reciprocal regulation of C/EBPα and inflammatory signaling. Blood. 2018;131(10):1081–93.

37. Xu X, Xiong X, Sun Y. The role of ribosomal proteins in the regulation of cell proliferation, tumorigenesis, and genomic integrity. Sci China Life Sci. 2016;59(7):656–72.

38. Land RH, Rayne AK, Vanderbeck AN, Barlowe TS, Manjunath S, Gross M, et al. The orphan nuclear receptor NR4A1 specifies a distinct subpopulation of quiescent myeloid-biased long-term HSCs. Stem Cells. 2015;33(1):278–88.

39. Schmidt MHH, Bicker F, Nikolic I, Meister J, Babuke T, Picuric S, et al. Epidermal growth factor-like domain 7 (EGFL7) modulates Notch signalling and affects neural stem cell renewal. Nat Cell Biol. 2009;11(7):873–80.

40. Richter A, Alexdottir MS, Magnus SH, Richter TR, Morikawa M, Zwijsen A, et al. EGFL7 Mediates BMP9-Induced Sprouting Angiogenesis of Endothelial Cells Derived from Human Embryonic Stem Cells. Stem Cell Reports. 2019;12(6):1250–9.

