## Supplemental Data for "Regulation of hematopoietic stem cell (HSC) proliferation by Epithelial Growth Factor Like-7 (EGFL7)"

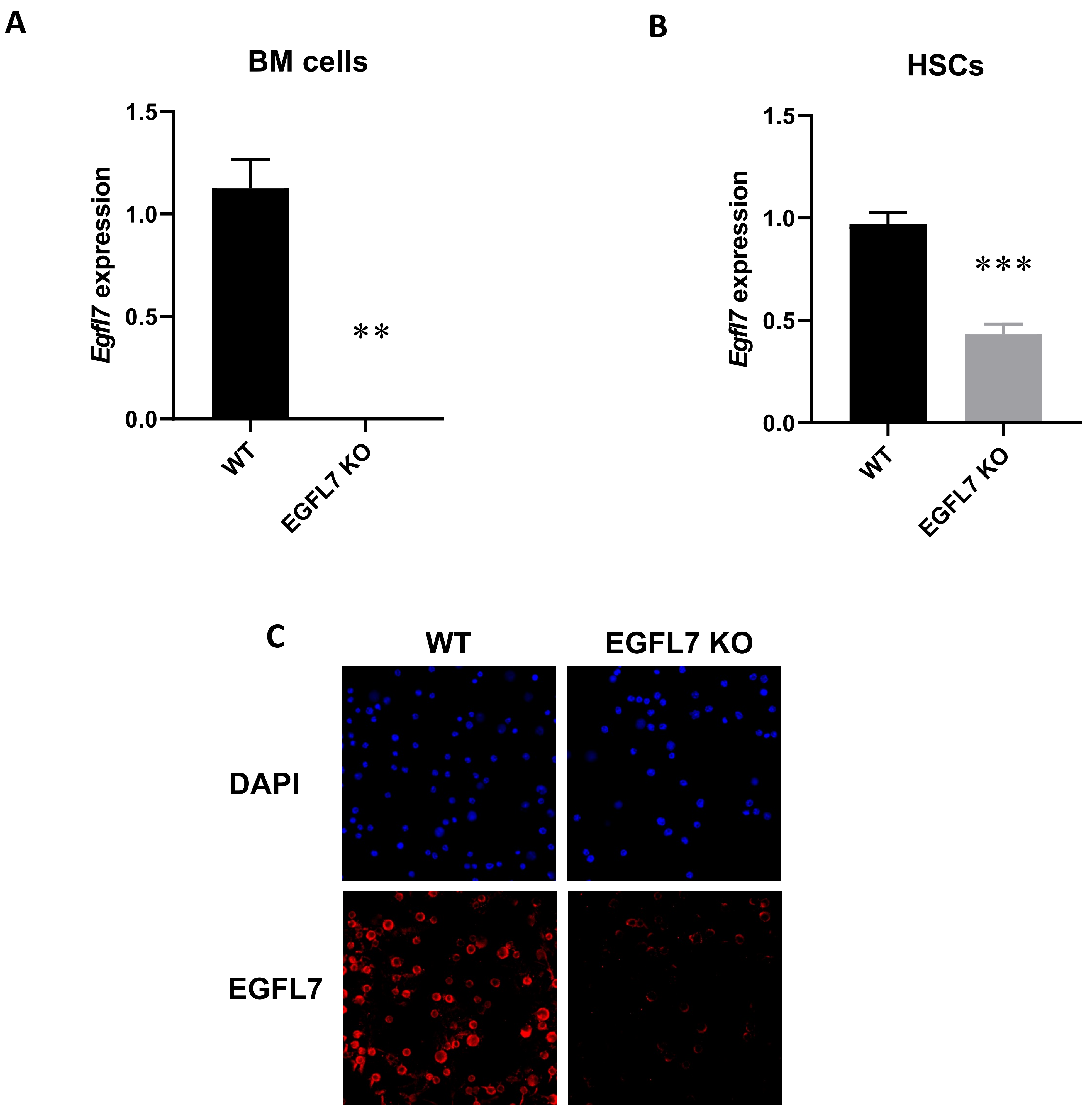
A
BM cells
Egfl7 expression
1.5
1.0
0.5
0.0
**
WT
EGFL7 KO
B
HSCs
Egfl7 expression
1.5
1.0
0.5
0.0
***
WT
EGFL7 KO
C
WT
EGFL7 KO
DAPI
EGFL7

**Figure S1**

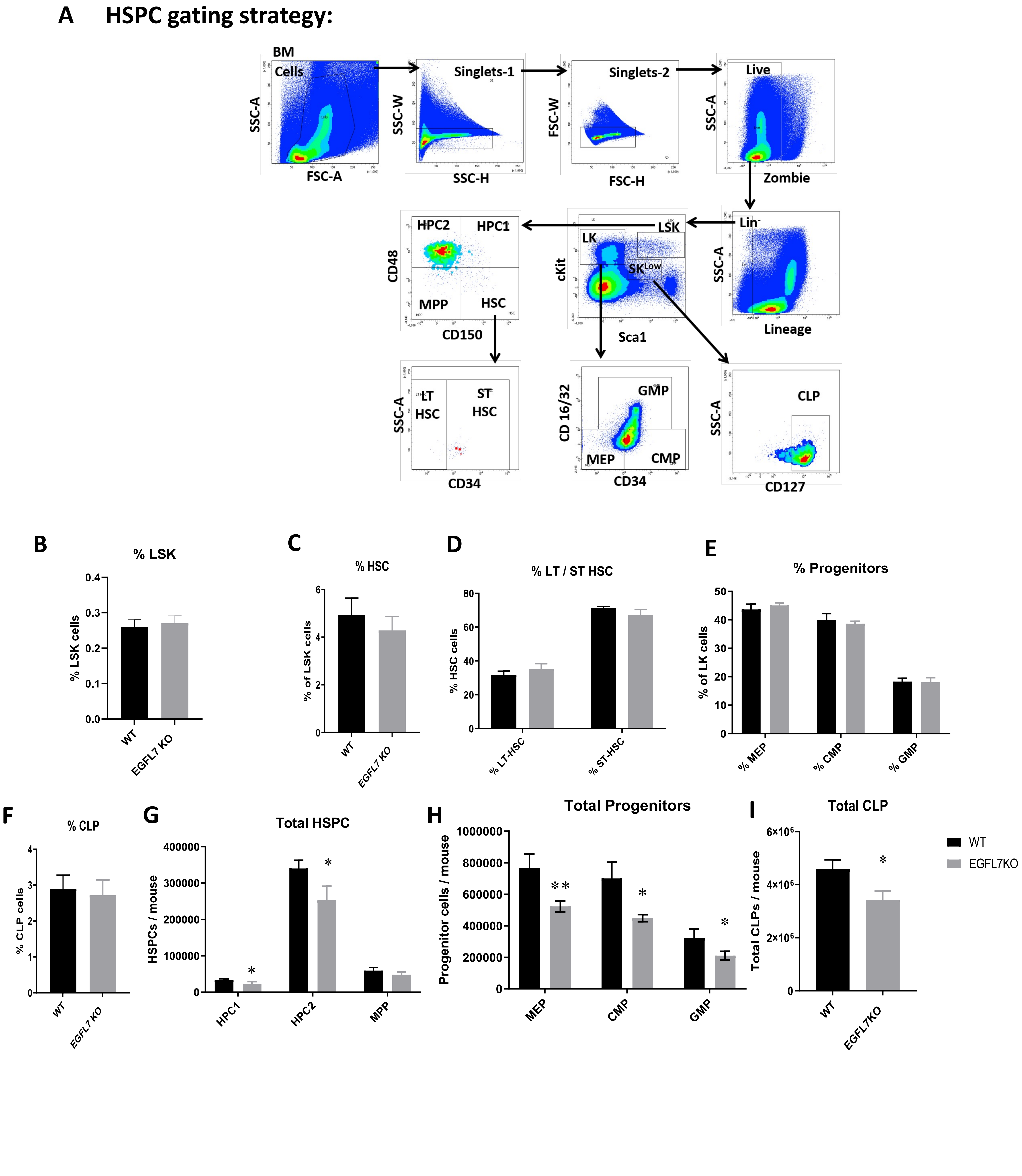
A
HSPC gating strategy:
BM
Cells
SSC-A
FSC-A
Singlets-1
SSC-W
SSC-H
Singlets-2
FSC-W
FSC-H
Live
SSC-A
Zombie
Lin-
SSC-A
Lineage
LK
LSK
SKLow
cKit
Sca1
HPC2
HPC1
MPP
HSC
CD48
CD150
LT HSC
ST HSC
SSC-A
CD34
GMP
MEP
CMP
CD 16/32
CD34
CLP
SSC-A
CD127
B
% LSK
% LSK cells
WT
EGFL7 KO
C
% HSC
% of LSK cells
WT
EGFL7 KO
D
% LT / ST HSC
% HSC cells
% LT-HSC
% ST-HSC
E
% Progenitors
% of LK cells
% MEP
% CMP
% GMP
F
% CLP
% CLP cells
WT
EGFL7 KO
G
Total HSPC
HSPCs / mouse
HPC1
HPC2
MPP
H
Total Progenitors
Progenitor cells / mouse
MEP
CMP
GMP
I
Total CLP
Total CLPs / mouse
WT
EGFL7KO
WT
EGFL7KO

Figure S2

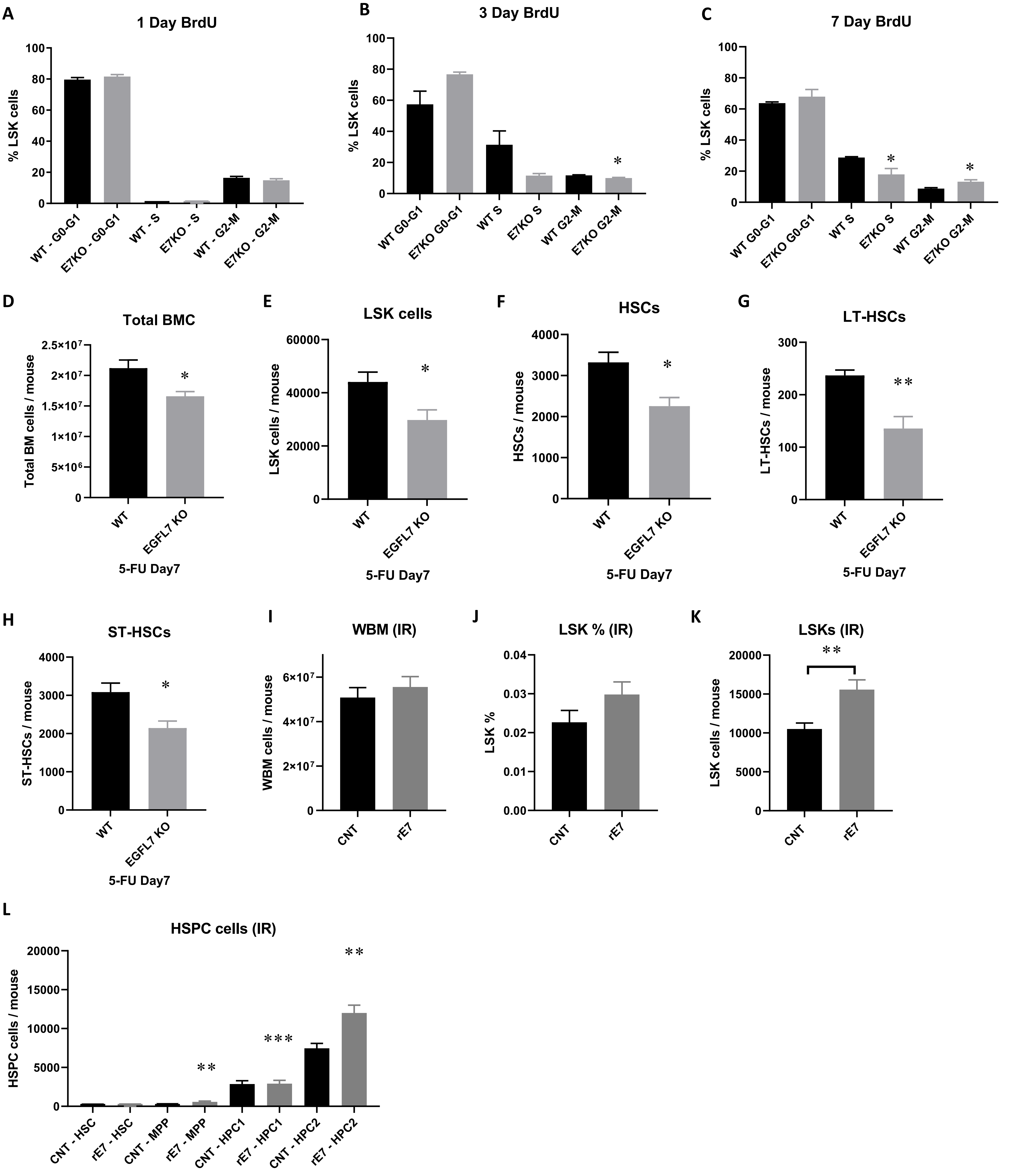
A
1 Day BrdU
% LSK cells
WT - G0-G1
E7KO - G0-G1
WT - S
E7KO - S
WT - G2-M
E7KO - G2-M
B
3 Day BrdU
% LSK cells
WT G0-G1
E7KO G0-G1
WT S
E7KO S
WT G2-M
E7KO G2-M
C
7 Day BrdU
% LSK cells
D
Total BMC
Total BM cells / mouse
WT
EGFL7 KO
5-FU Day7
E
LSK cells
LSK cells / mouse
F
HSCs
HSCs / mouse
G
LT-HSCs
LT-HSCs / mouse
H
ST-HSCs
ST-HSCs / mouse
I
WBM (IR)
WBM cells / mouse
CNT
rE7
J
LSK % (IR)
LSK %
K
LSKs (IR)
LSK cells / mouse
L
HSPC cells (IR)
HSPC cells / mouse
CNT - HSC
rE7 - HSC
CNT - MPP
rE7 - MPP
CNT - HPC1
rE7 - HPC1
CNT - HPC2
rE7 - HPC2

### Figure S3

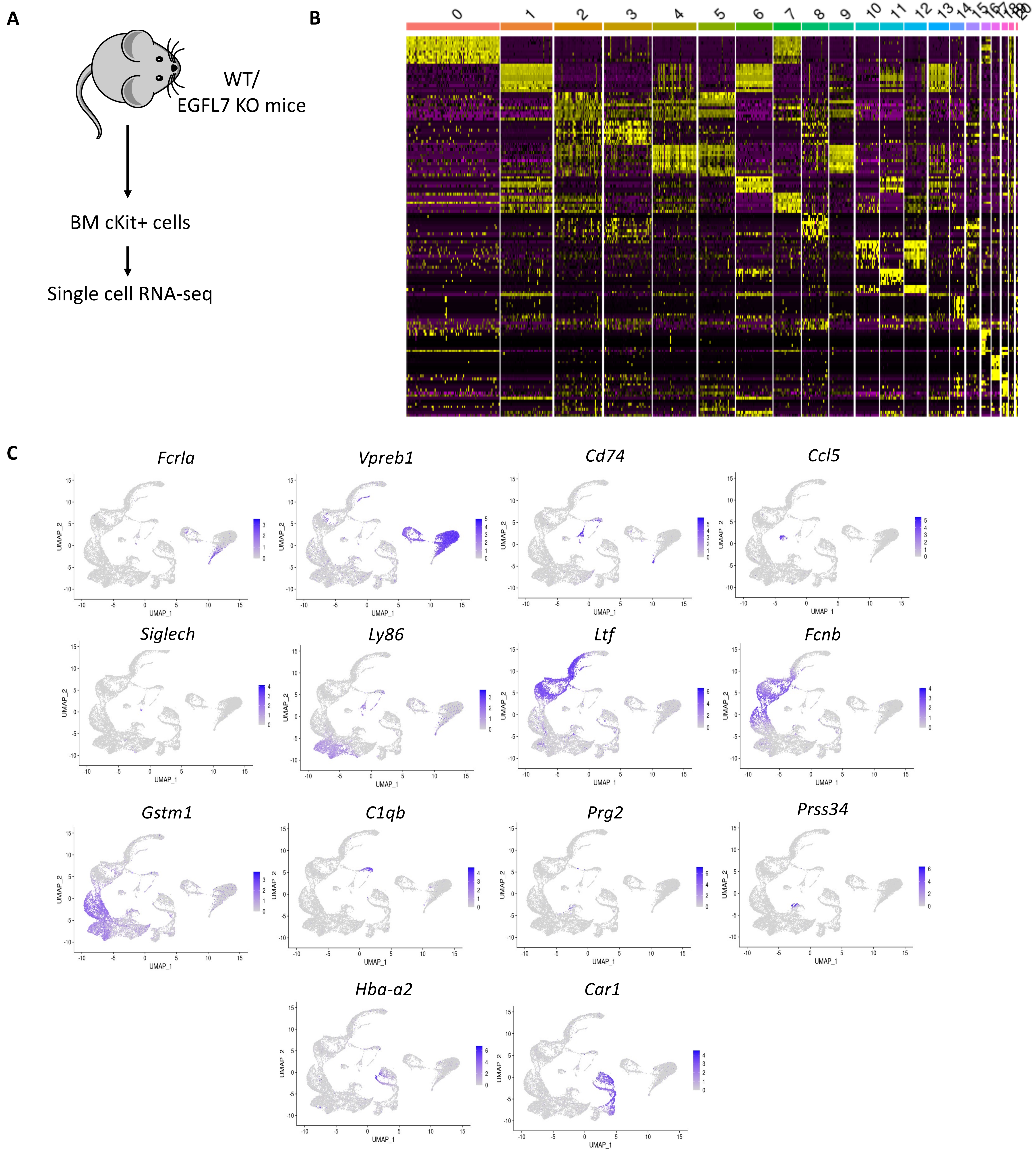
A
WT/
EGFL7 KO mice
BM cKit+ cells
Single cell RNA-seq
B
C
Fcrla
Vpreb1
Cd74
Ccl5
Siglech
Ly86
Ltf
Fcnb
Gstm1
C1qb
Prg2
Prss34
Hba-a2
Car1
UMAP_1
UMAP_2

### Figure S4

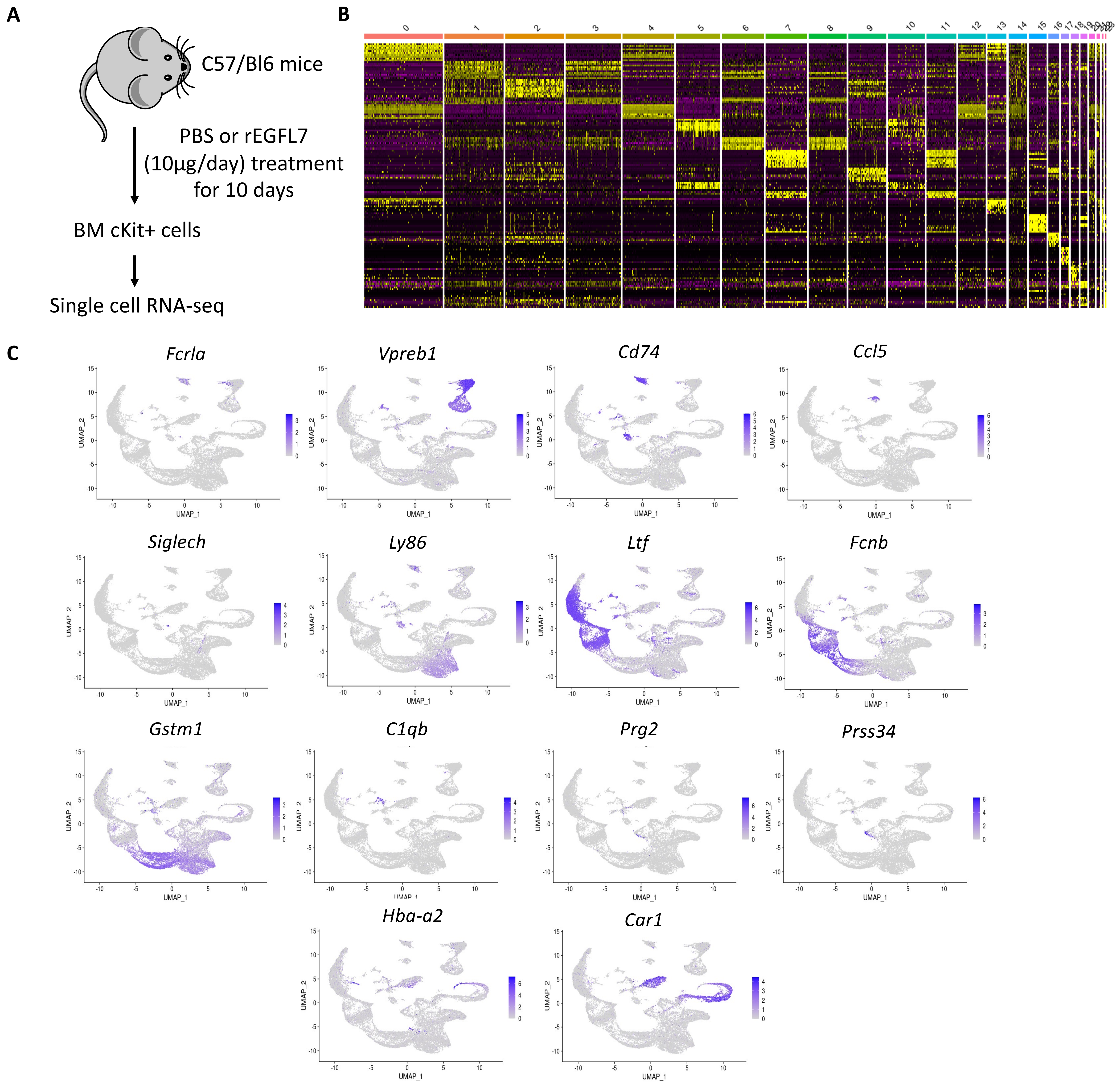
A
C57/Bl6 mice
PBS or rEGFL7
(10μg/day) treatment
for 10 days
BM cKit+ cells
Single cell RNA-seq
B
C
Fcrla
Vpreb1
Cd74
Ccl5
Siglech
Ly86
Ltf
Fcnb
Gstm1
C1qb
Prg2
Prss34
Hba-a2
Car1
UMAP_1
UMAP_2

### Figure S5

**SUPPLEMENTAL FIGURE LEGENDS**

**Supplemental Figure S1. EGFL7 expression in *Egfl7* KO mice.**

**(A)** BM cells and HSCs were sorted from 12-16 weeks old WT or *Egfl7* KO mice and *Egfl7* mRNA and EGFL7 protein expression were analyzed by qRT-PCR and immunofluorescence respectively. Relative expression of *Egfl7* in BM cells and **(B)** sorted HSCs from *Egfl7* KO mice compared to age-matched WT. Expression is relative to *Gapdh*. **(C)** Representative immunofluorescence image of BM mononuclear cells stained with EGFL7 (red) from *Egfl7* KO mice compared to WT. Nuclei are counterstained with DAPI. These data represent 3 or more experiments performed in triplicates with 3-6 mice per group. *P* = **<0.01, ***<0.001

**Supplemental Figure S2. *Egfl7* KO mice have normal BM HSPC frequency.**

**(A)** Representative gating strategy for identification of HSPC populations. BM of 12-16 weeks old *Egfl7* KO mice was analyzed for HSPC populations by flow cytometry. Age-matched WT mice were used as controls. **(B)** Percentage of LSK **(C)** percentage of HSCs, **(D)** percentage of LT and ST HSCs, **(E)** percentage of MEP, CMP, GMP, **(F)** percentage of CLP, **(G)** total number of HPC1, HPC2, MPP, **(H)** total number of MEP, CMP, GMP, and **(I)** total number of CLP in BM of *Egfl7* KO mice compared to WT. These data represent 3 or more experiments performed in triplicates with 3-6 mice per group. *P* = *<0.05, **<0.01, ***<0.001.

**Supplemental Figure S3. *Egfl7* loss delays cell cycle progression and impairs restoration of hematopoiesis post 5-FU and irradiatioin stress.**

*Egfl7* KO mice and age-matched WT controls were treated with a single injection (i.p) of BrdU (50 µg/g body weight). BrdU treatment continued through drinking water (0.8 mg/ml) until mice were sacrificed at 1-, 3-, 7-, and 30-days post BrdU injection. BM populations were analyzed for BrdU incorporation by flow cytometry. **(A)** Percentage of LSK cells in the $G_0$-$G_1$, S, and $G_2$- M cell cycle phases in *Egfl7* KO mice compared to control on day 1, **(B)** 3, and **(C)** 7 post BrdU treatment. **(D)** 12-16 weeks old WT or *Egfl7* KO mice injected with 5-FU (150 mg/kg, i.p.) were sacrificed 7 days post 5-FU injection and peripheral blood and BM was analyzed by an automated hematology analyzer and flow cytometry. Total number of BM cells, **(E)** number of BM LSK cells, **(F)** number of BM HSCs, **(G)** number of BM LT-HSCs, **(H)** number of BM ST-HSCs in WT or *Egfl7* KO mice on day 7 post 5-FU injection. **(I)** 16 weeks-old WT mice were irradiated with 5 Gy dose followed by treatment with rEGFL7 (rE7) or PBS (CNT) for 10 days, after which BM was analyzed by flow cytometry. Total number of BM cells, **(J)** percentage of BM LSKs, **(K)** total number of BM LSKs, and **(L)** total number of BM HSCs, MPP, HPC1, HPC2 cells in irradiated, rEGFL7-treated mice compared to control. These data are representative of 3 or more experiments with 4-6 mice per group. *P* = *<0.05, **<0.01, ***<0.001.

**Supplemental Figure S4. scRNA-sequencing of HSPCs from *Egfl7* KO mice.**

**(A)** Schematic representation of scRNA-seq experiment. BM from 12-16 weeks old *Egfl7* KO mice or age-matched WT controls was enriched for $cKit^+$ cells and submitted for scRNA-seq. **(B)** Heat map of top 25 genes expressed in each cluster. **(C)** UMAP of HSPCs colored by expression of cell-specific genes.

**Supplemental Figure S5. scRNA-sequencing of HSPCs from rEGFL7-treated mice.**

**(A)** Schematic representation of scRNA-seq experiment. 12-16 weeks old WT mice were treated with rEGFL7 or PBS (10µg/mouse, i.p) for 10 days after which mice were sacrificed, and BM cells were enriched for $cKit^+$ cells and submitted for scRNA-seq. **(B)** Heat map of top 25 genes expressed in each cluster. **(C)** UMAP of HSPCs colored by expression of cell-specific genes.

**SUPPLEMENTAL METHODS:**

**HSPC immunophenotyping and sorting of HSCs or LSK cells.**

HSPC immunophenotyping was performed as previously described (1). Briefly, 12-16 weeks old WT or *Egfl7* KO mice were sacrificed, and both pairs of tibia and femur were collected. BM was collected by crushing the bones in PBS + 2% FBS. BM cells were stained for various HSPC markers for 30 mins at 4°C. Following antibodies were used, mouse Lineage cocktail-PerCP/Cy5.5 (BD Pharmingen, Cat#51-9006964), anti-Ly6A/E-PE (BioLegend, Cat#108108), anti-CD117-APC/Cy7 (BioLegend, Cat#105826), anti-CD150-APC (BioLegend, Cat#115910), anti-CD48-PE/Cy7 (BioLegend, Cat#103424), anti CD34-FITC (BD Bioscience, Cat#553733), anti-CD16/32-eFlour450 (Invitrogen, Cat#48-0161-82), anti-CD127-PE/Dazzle594 (BioLegend, Cat#135031). Cells were washed with 1X DPBS and analyzed on BD LSR-II or Fortessa flow cytometer. Live cells were identified based on the exclusion of Zombie AquaTM dye (BioLegend, Cat# 423101).

For sorting HSCs or LSK cells, BM cells were layered on Histopaque®-1083 (Sigma, 10831-100ML) and centrifuged at 1600 rpm for 20 minutes at 25 °C. Buffy coat of mononuclear cells (MNCs) at the interface was collected and washed with 1X DPBS. Lineage-negative (Lin-) cells were enriched by staining with lineage cocktail (anti-CD5-biotin (BD Pharmingen, Cat#553019), anti-CD45R-biotin (BD Pharmingen, Cat#553086), anti-CD11b-biotin (BD Pharmingen, Cat#553309), anti-CD8a-biotin(BD Pharmingen, Cat#553029), anti-Gr1-biotin (BD Pharmingen, Cat#553125), anti-Ter119-biotin (BD Pharmingen, Cat#553672)) for 45 mins at 4°C. Cells were then washed and incubated with the Dynabeads$^{TM}$ Biotin Binder (Invitrogen, Cat#11047) for 30 mins. Lin- cells were collected as the flow through upon passage through magnetic separation column. Lin- cells were then stained for specific HSC or LSK markers listed below. Finally, HSCs or LSK cells were sort purified from the Lin- cells on BD- ARIA FACS machine.

Flow cytometry markers for identification of cells:

| No. | Cell type | Marker system for flow cytometry |
|---|---|---|
| | LT HSC | $Lin^-$ $Sca1^+$ $cKit^+$ $CD150^+$ $CD48^-$ $CD34^-$ |
| | ST HSC | $Lin^-$ $Sca1^+$ $cKit^+$ $CD150^+$ $CD48^-$ $CD34^+$ |
| | HSC | $Lin^-$ $Sca1^+$ $cKit^+$ $CD150^+$ $CD48^-$ |

| | | |
|---|---|---|
| | MPP | $Lin^- Sca1^+ cKit^+ CD150^- CD48^-$ |
| | HPC1 | $Lin^- Sca1^+ cKit^+ CD150^+ CD48^+$ |
| | HPC2 | $Lin^- Sca1^+ cKit^+ CD150^- CD48^+$ |
| | LSK | $Lin^- Sca1^+ cKit^+$ |
| | LK | $Lin^- Sca1^- cKit^+$ |
| | MEP | $Lin^- Sca1^- cKit^+ CD34^- CD16/32^-$ |
| | CMP | $Lin^- Sca1^- cKit^+ CD34^+ CD16/32^-$ |
| | GMP | $Lin^- Sca1^- cKit^+ CD34^+ CD16/32^+$ |
| | CLP | $Lin^- Sca1^{low} cKit^{low} CD127^+$ |

**LTC-IC assay:**

LTC-IC assays were performed by plating ($5X10^4$ cells/well) BM cells on primary murine MSC feeder layer. The culture was maintained at 37°C in 5% $CO_2$ with ≥ 95% humidity for 4 weeks, with weekly half media change. After the incubation period, LTC-IC cultures were harvested, and a CFU-C assay was performed in MethoCult$^{TM}$ M3434 (Stem Cell Technologies). Cultures were then incubated for twelve to fourteen days at 37°C in 5% $CO_2$ with ≥ 95% humidity. The total number of colonies per dish was counted. The total number of LTC-IC-derived CFU was calculated using seeded cell number at day 0, harvested cell number at week 4, and seeded cell number at week 4.

**Transplantation studies:**

For Non-competitive transplantation experiments, WT or *Egfl7* KO C57BL/6J (CD45.2) mice of 12-16 weeks age were used as the donors. WT B6.SJL-Ptprca Pep3b/BoyJ (CD45.1) mice at the age of 8-10 weeks were used as the recipients. $2X10^6$ Donor $CD45.2^+$ BM cells were mixed with $1X10^5$ freshly isolated $CD45.1^+$ MNCs (rescue cells) and i.v. transplanted into lethally irradiated (10 Gy, x-irradiation, given in two split doses of 5 Gy at 3 hours apart) BoyJ (CD45.1) mice. The levels of chimerism and differentiated cell formation was estimated every 4 weeks up to 16 weeks post-transplantation.

For competitive transplantation experiments, WT or *Egfl7* KO C57BL/6J (CD45.2) mice of 12-16 weeks age were used as the donors. WT B6.SJL-Ptprca Pep3b/BoyJ (CD45.1) mice at the age of 8-10 weeks were used as the recipients. $2X10^6$ Donor $CD45.2^+$ BM cells were mixed with $2X10^6$ freshly isolated $CD45.1^+$ MNCs (rescue cells) and i.v. transplanted into lethally irradiated (10 Gy, x-irradiation, given in two split doses of 5 Gy at 3 hours apart) BoyJ (CD45.1) mice. The levels of chimerism and differentiated cell formation was estimated every 4 weeks up to 16 weeks post-transplantation.

The engraftment potential was examined by performing BM analysis of the recipient mice at 16 weeks post-transplantation. For secondary transplantations, engrafted CD45.2$^{+}$ cells were sort-purified from the primary recipients' bone marrow and transplanted into secondary recipients like the primary transplantation protocol mentioned earlier.

**Sublethal irradiation:**

Mice were irradiated with a single dose of 5.0 Gy (X-ray irradiation) and BM was analyzed on day 10 post-irradiation.

**Single-cell RNA sequencing:**

Single-cell RNA sequencing (scRNAseq) was performed using 10X Genomic's 3' gene expression kit (v3.1, dual indexed). We harvested HSPCs from *Egfl7* KO and WT mice BM, as well as WT mice treated for 10 days with 10ug of rEGFL7 and PBS controls. BM was isolated by crushing pooled tibias and femurs from each group. Mononuclear cells were obtained by ficol gradient centrifugation and cKit+ cells were isolated using MACS (Milteny). All cells were resuspended at 1000cells/µl and sequenced. Cell suspensions were loaded into a 10xGenomics Chromium device for microfluidic-based partitioning and capture of single cells. The manufacturer's instructions were followed for reverse transcription, cDNA amplification, and library preparation according to the v.3.1 3'-single-cell RNA-seq protocol. Sequencing was performed on an Illumina NovaSeq 6000 instrument to generate paired-end sequencing reads, and FASTQ files were generated using Illumina bcl2fastq software. 10x Genomics CellRanger software suite was used following the default parameters, to perform data pre-processing including alignment, filtering, barcode counting, and UMI counting. Downstream analyses were performed using Seurat V3 for R (2). Briefly, the feature-barcode matrices were log-normalized, and top variable genes across single cells were identified using the FindVariableGenes function. Dimensionality reduction was performed using principal component analysis (PCA) and then the distance matrix was organized into a K-nearest neighbor graph (KNN), partitioned into clusters using the Louvain algorithm, and clusters were visualized on a UMAP or t-SNE plot. Top differentially expressed genes for each cluster were computed using the FindMarkers function, and cell types were annotated based on the expression of known markers

**Immunofluorescence Microscopy (IFC):**

We sort purified HSC and LK cells from BM of 12-16 weeks-old mice using flow cytometry. These cells were allowed to adhere to fibronectin-coated slides for 1 hour. The cells were then both fixed using 1% PFA and permeabilized using 0.1% Triton X for 10 mins at room temperature. Cells were then stained with primary antibodies against EGFL7 (rabbit anti-mouse IgG; LS Bio), Ki67 (rat anti-mouse IgG; Invitrogen) followed by secondary antibodies - Alexa Fluor 594 (donkey anti-rat IgG; Fisher Scientific) and Alexa Fluor 647 (donkey anti-rabbit IgG; Abcam) respectively. We counterstained the nuclei with 4,6 diamidino-2-phenylindole (DAPI) before imaging on a confocal microscope.

**Real-time quantitative PCR:**

Total RNA was isolated using RNeasy Plus Micro kit (Qiagen) according to manufacturer's instructions. cDNA was synthesized using 500 ng of RNA using Radiant cDNA Synthesis Kit (Radiant Molecular Tools). The expressions of respective genes were analyzed using SuperScript IV VILO Master Mix (Thermo Fisher Scientific). Values were then normalized to the expression of GAPDH for each sample. Expression of the following genes were analyzed using TaqMan® Gene Expression Assays (Thermo Fisher Scientific):

1. Egfl7 (Assay ID - Mm00618006_g1)
2. Egr1 (Assay ID - Mm00656724_m1)
3. Klf2 (Assay ID - Mm00500486_g1)
4. Klf4 (Assay ID - Mm00516104_m1)
5. Mid1 (Assay ID - Mm04933165_m1)
6. Nfkbiz (Assay ID - Mm00600522_m1)
7. Nr4a1 (Assay ID - Mm01300401_m1)
8. Nr4a2 (Assay ID - Mm01300401_m1)
9. S100a8 (Assay ID - Mm00496696_g1)
10. S100a9 (Assay ID - Mm00443060_m1)
11. Vegfa (Assay ID - Mm00437306_m1)
12. ActB (Assay ID - Mm00607939_s1)

**Statistical Analysis:**

Data analysis on the examined groups was performed using Graph Pad Prism 8.0 software and comparative analysis using a t-test. Data are reported as mean values ± standard deviation. Statistical significance is as follows: *$p<0.05$, **$p<0.01$, ***$p<0.001$.

**REFERENCES**

1. Oguro H, Ding L, Morrison SJ. SLAM family markers resolve functionally distinct subpopulations of hematopoietic stem cells and multipotent progenitors. Cell Stem Cell. 2013;13(1):102-16.

2. Stuart T, Butler A, Hoffman P, Hafemeister C, Papalexi E, Mauck WM, et al. Comprehensive Integration of Single-Cell Data. Cell. 2019;177(7):1888-902.e21.
